# Mechanism and spectrum of inhibition of a 4’-cyano modified nucleotide analog against diverse RNA polymerases of prototypic respiratory RNA viruses

**DOI:** 10.1101/2024.04.22.590607

**Authors:** Calvin J. Gordon, Simon M. Walker, Egor P. Tchesnokov, Dana Kocincova, Jared Pitts, Dustin S. Siegel, Jason K. Perry, Joy Y. Feng, John P. Bilello, Matthias Götte

## Abstract

The development of safe and effective broad-spectrum antivirals that target the replication machinery of respiratory viruses is of high priority in pandemic preparedness programs. Here, we studied the mechanism of action of a newly discovered nucleotide analog against diverse RNA-dependent RNA polymerases (RdRp) of prototypic respiratory viruses. GS-646939 is the active 5′-triphosphate (TP) metabolite of a 4ʹ-cyano modified *C*-adenosine analog phosphoramidate prodrug GS-7682. Enzyme kinetics show that the RdRps of human rhinovirus type 16 (HRV-16) and enterovirus 71 (EV-71) incorporate GS-646939 with unprecedented selectivity; GS-646939 is incorporated 20-50-fold more efficiently than its natural ATP counterpart. The RdRp complex of respiratory syncytial virus (RSV) and human metapneumovirus (HMPV) incorporate GS-646939 and ATP with similar efficiency. In contrast, influenza B RdRp shows a clear preference for ATP and human mitochondrial RNA polymerase (h-mtRNAP) does not show significant incorporation of GS-646939. Once incorporated into the nascent RNA strand, GS-646939 acts as a chain-terminator although higher NTP concentrations can partially overcome inhibition for some polymerases. Modeling and biochemical data suggest that the 4ʹ-modification inhibits RdRp translocation. Comparative studies with GS-443902, the active triphosphate form of the 1′-cyano modified prodrugs remdesivir and obeldesivir, reveal not only different mechanisms of inhibition, but also differences in the spectrum of inhibition of viral polymerases. In conclusion, 1ʹ-cyano and 4ʹ-cyano modifications of nucleotide analogs provide complementary strategies to target the polymerase of several families of respiratory RNA viruses.

## INTRODUCTION

Respiratory RNA viruses represent a substantial public health burden worldwide. Facile transmission from person to person can cause outbreaks, epidemics, or pandemics. Severe acute respiratory syndrome coronavirus-2 (SARS-CoV-2), the causative agent of coronavirus disease 2019 (COVID-19)(1,2), is a most recent example. Other prominent examples include members of the *Coronaviridae* such as SARS-CoV and Middle East respiratory syndrome coronavirus (MERS-CoV); the *Picornaviridae*, e.g. human rhinovirus (HRV); the *Pneumoviridae*, e.g. respiratory syncytial virus (RSV); the *Paramyxoviridae*, e.g. human parainfluenza viruses (HPIV); and influenza viruses of the *Orthomyxoviridae*. Infection with these pathogens is associated with diverse disease outcomes, from asymptomatic or mild sequelae to viral bronchiolitis and pneumonia. High rates of hospitalizations and mortality from viral respiratory infections are of particular concern in children, older adults, and those with chronic airway inflammatory diseases, such as asthma (3–8).

The development of effective medical countermeasures is challenging due to the diverse nature of the aforementioned viruses that cover positive-sense RNA viruses (coronaviruses and picornaviruses), non-segmented negative-sense RNA viruses (pneumoviruses and paramyxoviruses) and segmented negative-sense RNA viruses (orthomyxoviruses). A common target for pharmaceutical intervention strategies is the viral RNA-dependent RNA polymerase (RdRp), which is required for viral genome replication. Although the structural details of these enzymes differ across virus families and to a lesser degree from virus species to species, the active site is relatively conserved to accommodate nucleoside 5′-triphosphate (NTP) substrates. The development of antiviral nucleoside or nucleotide analogs is therefore a logical strategy to identify therapeutics with potential for broad-spectrum antiviral activity. Once incorporated into the growing RNA chain, the nucleotide analog can cause inhibition of RNA synthesis. The detailed mechanism of action depends on both the nature of the inhibitor and the nature of the polymerase.

Remdesivir (RDV) is a 1ʹ-cyano modified *C*-adenosine monophosphate prodrug derived from the parent nucleoside GS-441524 (Fig. 1*A*) (9). RDV was the first antiviral drug to receive approval from the US Food and Drug Administration (FDA) for the treatment of COVID-19 (10). *In vitro*, RDV and GS-441524 exhibit broad-spectrum antiviral activity against respiratory viruses, including coronaviruses (SARS-CoV-2, SARS-CoV, and MERS-CoV) (11–15), picornaviruses (HRV-16, enterovirus D68 [EV-D68]) (16,17), pneumoviruses (RSV, human metapneumovirus [HMPV]) (18,19), and human paramyxoviruses (HPIV) (19,20). Antiviral activity is not observed against several segmented negative-sense RNA viruses, including Lassa virus (LASV), Crimean-Congo hemorrhagic fever virus (CCHFV), and influenza viruses (17,19,20). The 5ʹ ester prodrug of GS-441524, obeldesivir (ODV), is metabolized pre-systemically to GS-441524 and subsequently to GS-443902, and therefore shares the same antiviral profile as RDV and its parent nucleoside (21). ODV was recently evaluated in phase 3 clinical trials for COVID-19 and was efficacious against filoviruses in a post-exposure prophylaxis non-human primate model (22).

**Figure 1:**
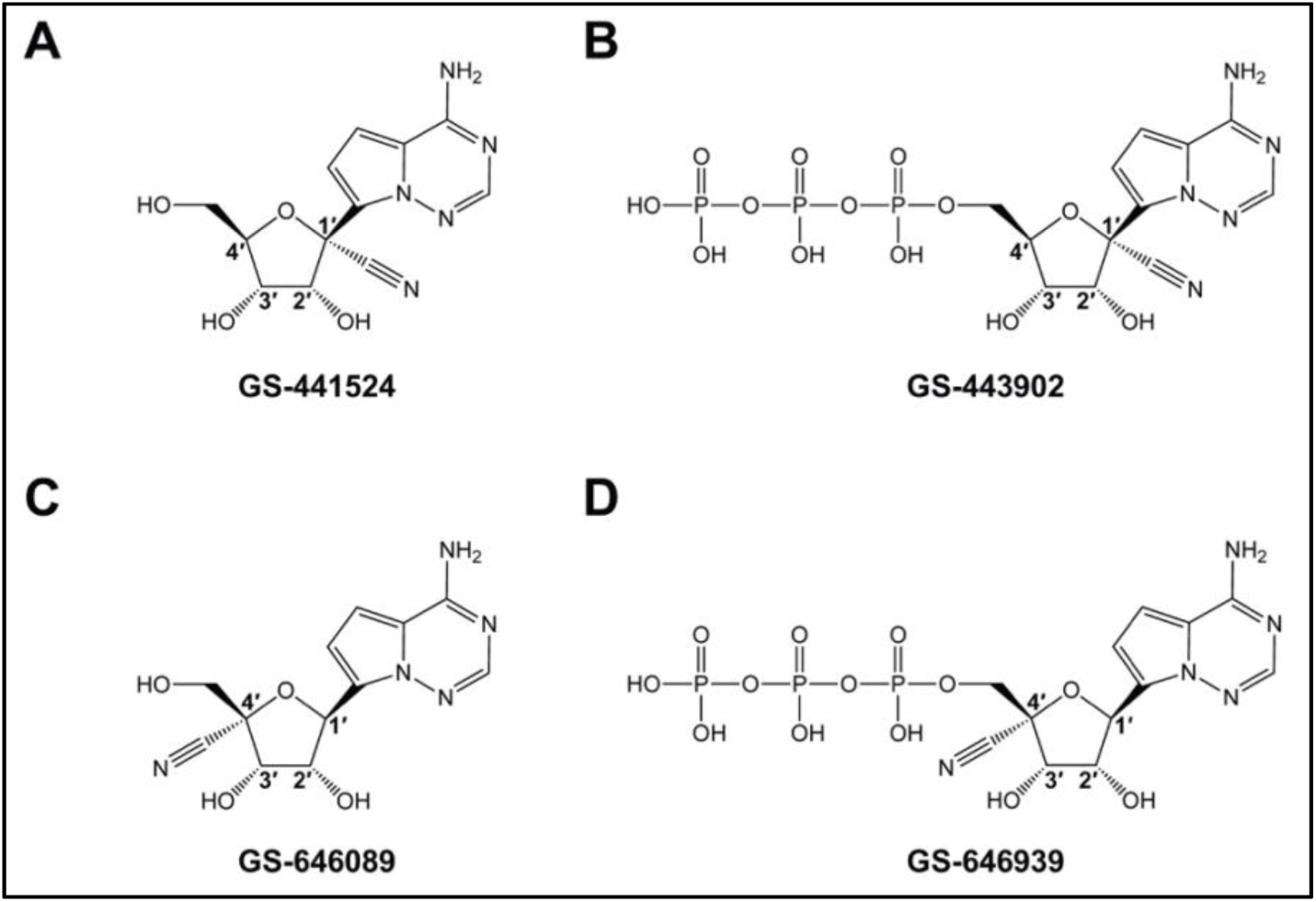
Chemical structures of GS-441524 (***A***), GS-443902 (***B***), GS-646089 (***C***), and GS-646939 (***D***).

Key biochemical attributes of RDV and GS-441524 that enable potent inhibition of the SARS-CoV-2 RdRp complex have been identified. Their active 5′-triphosphate metabolite, herein referred to as GS-443902 (Fig. 1*B*), outcompetes its natural counterpart adenosine triphosphate (ATP) 2-to 3-fold (23–25). Incorporation into the growing RNA chain at position “i” results in delayed chain-termination at position “i+3”. A steric clash between the 1ʹ-cyano group of GS-443902 and the conserved Ser-861 in the RdRp causes inhibition of primer extension (26,27). Higher NTP concentrations can overcome delayed-chain termination at position “i+3”, potentially yielding complete copies of the viral genome with embedded GS-443902 residues (26–31). When used as templates, incorporation of the complementary UTP is likewise inhibited. This template-dependent inhibition is more effective, but higher NTP concentrations can overcome this obstacle as well (27).

Here we compare the 1ʹ-cyano modified *C*-adenosine with a newly discovered 4ʹ-cyano modified *C*-adenosine. Derived from the nucleoside GS-646089 (Fig. 1*C*), the monophosphate prodrug GS-7682 is associated with a broad-spectrum of antiviral activities (32). Viruses that belong to the *Picorna-* and *Pneumoviridae* families were most sensitive to GS-7682 treatment in cell culture. The 50% effective concentration (EC_50_) of GS-7682 was less than 100 nM against several picornaviruses, including HRV-16 (32). Further, GS-7682 was potent against HMPV and RSV with EC_50_ values of 210 ± 50 and 3-46 nM (various assays), respectively. However, GS-7682 demonstrated limited antiviral activity against corona- and orthomyxoviruses (32). In this study, we compared the biochemical properties of the active 5′-trisphosphate form of GS-7682, referred to as GS-646939 (Fig. 1*D*), with GS-443902 against an array of RdRp enzymes representing multiple families of respiratory viruses. Highly efficient rates of GS-646939 incorporation are seen with picornaviruses HRV-16 and EV-71 RdRp and, to a lesser extent, with pneumoviruses RSV and HMPV RdRp. In contrast to GS-443902, inhibition with GS-646939 is based on immediate chain-termination at position “i”. Overall, the biochemical evaluation of the two nucleotide analogs against their targets provides mechanistic detail for the observed antiviral effects.

## RESULTS

### Experimental strategy

The main objective of this study was to elucidate the mechanism of action of GS-646939 using purified RdRp enzymes from prototypic respiratory RNA viruses of the *Coronaviridae, Picornaviridae*, *Pneumoviridae*, *Paramyxoviridae* and *Orthomyxoviridae* (Table 1). The prototypic pathogen approach is based on research efforts with selected viruses that represent specific families (33). To strengthen this approach, we expressed the RdRp from two prototypic species within each of the five families mentioned above. SARS-CoV-2 and MERS-CoV were selected to represent coronaviruses, HRV-16 and EV-71 for picornaviruses, RSV and HMPV for pneumoviruses, and HPIV-3 and PIV-5 for paramyxoviruses. Orthomyxoviruses were represented by influenza B virus. The RdRp from Lassa virus (LASV) a segmented negative-sense RNA virus (like influenza) which belongs to the *Arenaviridae* family, was also evaluated. Human mitochondrial RNA polymerase (h-mtRNAP) was utilized to assess potential off-target effects. The comprehensive approach was designed to identify common biochemical properties that are associated with broad-spectrum antiviral activity. Active RdRp enzymes used in this study exist either as monomers, dimers or multimeric complexes (Table 1). The coronaviruses SARS-CoV-2 and MERS-CoV form complexes with non-structural protein 12 (Nsp12), which contains the RdRp active site, and essential cofactors Nsp7 and Nsp8 (25,34,35). Picornavirus RdRps from HRV-16 and EV-71, also referred to as 3D^pol^, are monomeric and do not require additional cofactors (36,37). The replication complex of the pneumoviruses RSV and HMPV, as well as paramyxoviruses HPIV-3 and PIV-5, consists of the large (L) protein, containing the RdRp active site, and a phosphoprotein (P) (38,39). PIV-5, or canine parainfluenza virus, was included because the structure of the RdRp complex is available (40) (Table 1). Active recombinant HPIV-3 RdRp has not yet been reported. Influenza viruses assemble a heterotrimeric complex composed of PA (cap-snatching endonuclease subunit), PB1 (RdRp subunit), and PB2 (cap-binding subunit) (41). The LASV L protein is a dynamic monomer functionally similar to the trimeric influenza replication complex, possessing an N-terminus endonuclease, RdRp core, and C-terminus cap-binding domain (42). Finally, h-mtRNAP is a single subunit enzyme responsible for the transcription of the mitochondrial genome (43). In this study, we combined enzymatic assays with structural modeling to provide insight into the requirements for nucleotide incorporation and inhibition of RNA synthesis, respectively, with GS-443902 serving as a benchmark.

**Table 1:**
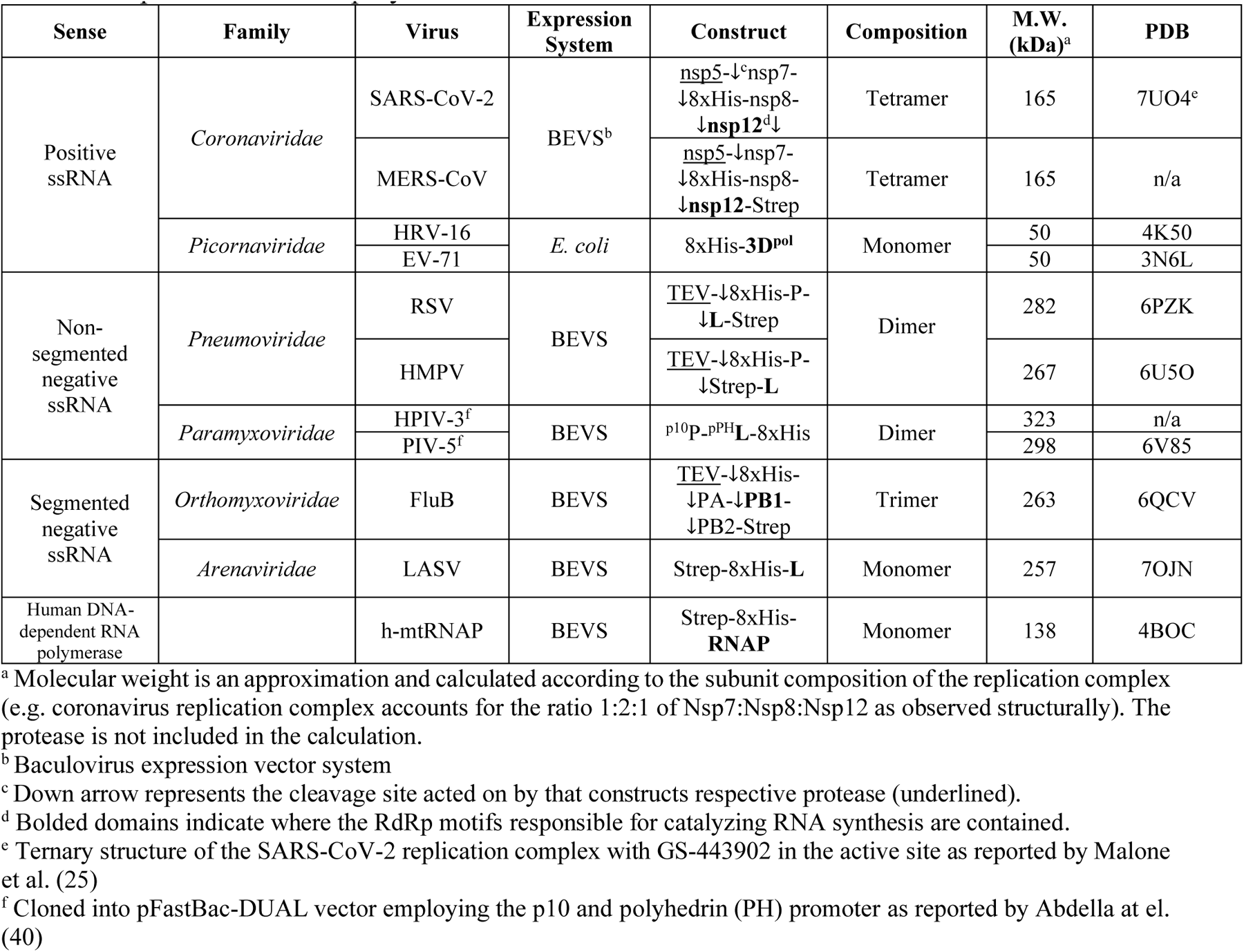
Expression of RNA polymerases.

### Selective incorporation of GS-443902 and GS-646939 by representative RdRp enzymes

We compared nucleotide incorporation efficiency of the 1ʹ-cyano GS-443902 (Table 2) with the 4ʹ-cyano GS-646939 modified NTP (Table 3). Enzyme reactions were monitored in gel-based assays using model primer/templates (Fig. S1) (26,44–47). Steady-state kinetic parameters V_max_/*K*_m_ for ATP incorporation over V_max_/*K*_m_ for the nucleotide analog provide a measure of the efficiency of nucleotide incorporation in relation to its natural counterpart. The numerical value is unitless and defines the selectivity for a given nucleotide analog. Selectivity values below 1 indicate that the nucleotide analog is more efficiently incorporated than ATP. Previous studies revealed selectivity values of ∼0.3 for GS-443902 incorporation by RdRp complexes of coronaviruses SARS-CoV-2, SARS-CoV, and MERS-CoV (Table 2) (23,26,27,44). HRV-16 and EV-71 RdRp show selectivity values around 1 for GS-443902, indicating that picornavirus enzymes incorporate this nucleotide analog and ATP with similar efficiency. RdRp complexes of pneumoviruses RSV and HMPV show higher selectivity values of 2.7 and 6.7, respectively, signifying a preference for the natural ATP nucleotide. A similar observation was made with paramyxovirus enzyme complexes of HPIV-3 (7.2) and PIV-5 (8.3). Incorporation of GS-443902 by influenza B and LASV RdRp is even more limited, with selectivity values of 70 and 19, respectively. GS-443902 incorporation by h-mtRNAP is highly inefficient, with a selectivity value of ∼500 (46).

**Table 2:** Selective incorporation of GS-443902, a 1ʹ-cyano purine NTP analog, by selected RdRp enzymes.

| RNA Sense | Family (-viridae) | Virus | ATP |  |  | GS-443902 |  |  | Selectivity<sup>f</sup> (fold) |
| --- | --- | --- | --- | --- | --- | --- | --- | --- | --- |
| | | | $V_{\max}^a$<br>(product fraction) | $K_m^b$ (μM) | $V_{\max}/K_m$ | $V_{\max}$<br>(product fraction) | $K_m$ (μM) | $V_{\max}/K_m$ | |
| Positive ssRNA | Corona- | **SARS-CoV-2** |  |  |  |  |  |  | **0.30<sup>1</sup>** |
|  |  | P.R.<sup>c</sup> | 0.75<sup>d</sup> | 0.03 | 25 | 0.74 | 0.0089 | 83 |  |
|  |  | ±<sup>e</sup> | 0.019 | 0.003 |  | 0.023 | 0.0020 |  |  |
|  |  | **SARS-CoV** |  |  |  |  |  |  | **0.34<sup>1</sup>** |
|  |  | P.R. | 0.73 | 0.03 | 24 | 0.70 | 0.010 | 70 |  |
|  |  | ± | 0.017 | 0.003 |  | 0.015 | 0.0008 |  |  |
|  |  | **MERS-CoV** |  |  |  |  |  |  | **0.35<sup>1</sup>** |
|  |  | P.R. | 0.47 | 0.017 | 28 | 0.50 | 0.0063 | 79 |  |
|  |  | ± | 0.011 | 0.019 |  | 0.012 | 0.0006 |  |  |
|  | Picorna- | **HRV-16** |  |  |  |  |  |  | **1.03** |
|  |  |  | 0.90 | 4.52 | 0.20 | 0.96 | 4.96 | 0.19 |  |
|  |  | ± | 0.017 | 0.31 |  | 0.015 | 0.27 |  |  |
|  |  | **EV-71** |  |  |  |  |  |  | **0.93** |
|  |  |  | 0.96 | 0.96 | 1.0 | 0.93 | 0.86 | 1.08 |  |
|  |  | ± | 0.021 | 0.066 |  | 0.027 | 0.081 |  |  |
| Nonsegmented negative ssRNA | Pneumo- | **RSV** |  |  |  |  |  |  | **2.73<sup>2</sup>** |
|  |  | P.R. | 0.76 | 0.17 | 4.5 | 0.82 | 0.50 | 1.6 |  |
|  |  | ± | 0.022 | 0.023 |  | 0.027 | 0.089 |  |  |
|  |  | **HMPV** |  |  |  |  |  |  | **6.71** |
|  |  |  | 0.60 | 6.40 | 0.094 | 0.57 | 42.19 | 0.014 |  |
|  |  | ± | 0.024 | 0.98 |  | 0.045 | 7.79 |  |  |
|  | Paramyxo- | **HPIV-3** |  |  |  |  |  |  | **7.20** |
|  |  |  | 0.63 | 0.061 | 10.3 | 0.63 | 0.44 | 1.43 |  |
|  |  | ± | 0.025 | 0.015 |  | 0.010 | 0.040 |  |  |
|  |  | **PIV-5** |  |  |  |  |  |  | **8.31** |
|  |  |  | 0.74 | 1.5 | 0.49 | 0.77 | 13 | 0.059 |  |
|  |  | ± | 0.026 | 0.25 |  | 0.042 | 2.3 |  |  |
| Segmented Negative ssRNA | Orthomyxo- | **FluB** |  |  |  |  |  |  | **70<sup>1</sup>** |
|  |  | P.R. | 0.73 | 0.23 | 3.2 | 0.45 | 9.7 | 0.046 |  |
|  |  | ± | 0.014 | 0.020 |  | 0.0064 | 0.42 |  |  |
|  | Arena- | **LASV** |  |  |  |  |  |  | **19<sup>1</sup>** |
|  |  | P.R. | 0.57 | 0.11 | 5.2 | 0.35 | 1.3 | 0.27 |  |
|  |  | ± | 0.032 | 0.020 |  | 0.016 | 0.18 |  |  |
| Human DNA-dependent RNA polymerase |  | **h-mtRNAP** |  |  |  |  |  |  | **503<sup>2</sup>** |
|  |  | P.R. | 0.98 | 0.050 | 19.6 | 0.81 | 21 | 0.039 |  |
|  |  | ± | 0.018 | 0.0037 |  | 0.013 | 0.096 |  |  |

<sup>a</sup> $V_{\max}$ is a Michaelis–Menten parameter reflecting the maximal velocity of nucleotide incorporation.
<sup>b</sup> $K_m$ is a Michaelis–Menten parameter reflecting the concentration of the nucleotide substrate at which the velocity of nucleotide incorporation is half of $V_{\max}$
<sup>c</sup> Previously reported
<sup>d</sup> All reported values have been calculated based on an 8–data point experiment repeated at least 3 times (n=3).
<sup>e</sup> Standard error associated with the fit.
<sup>f</sup> Selectivity of a viral RNA polymerase for a nucleotide substrate analog is calculated as the ratio of the $V_{\max}/K_m$ values for ATP and GS-443902 analog, respectively.
<sup>1</sup> Data from Gordon et al. (44)
<sup>2</sup> Data from Tchesnokov et al. (46)

**Table 3:**
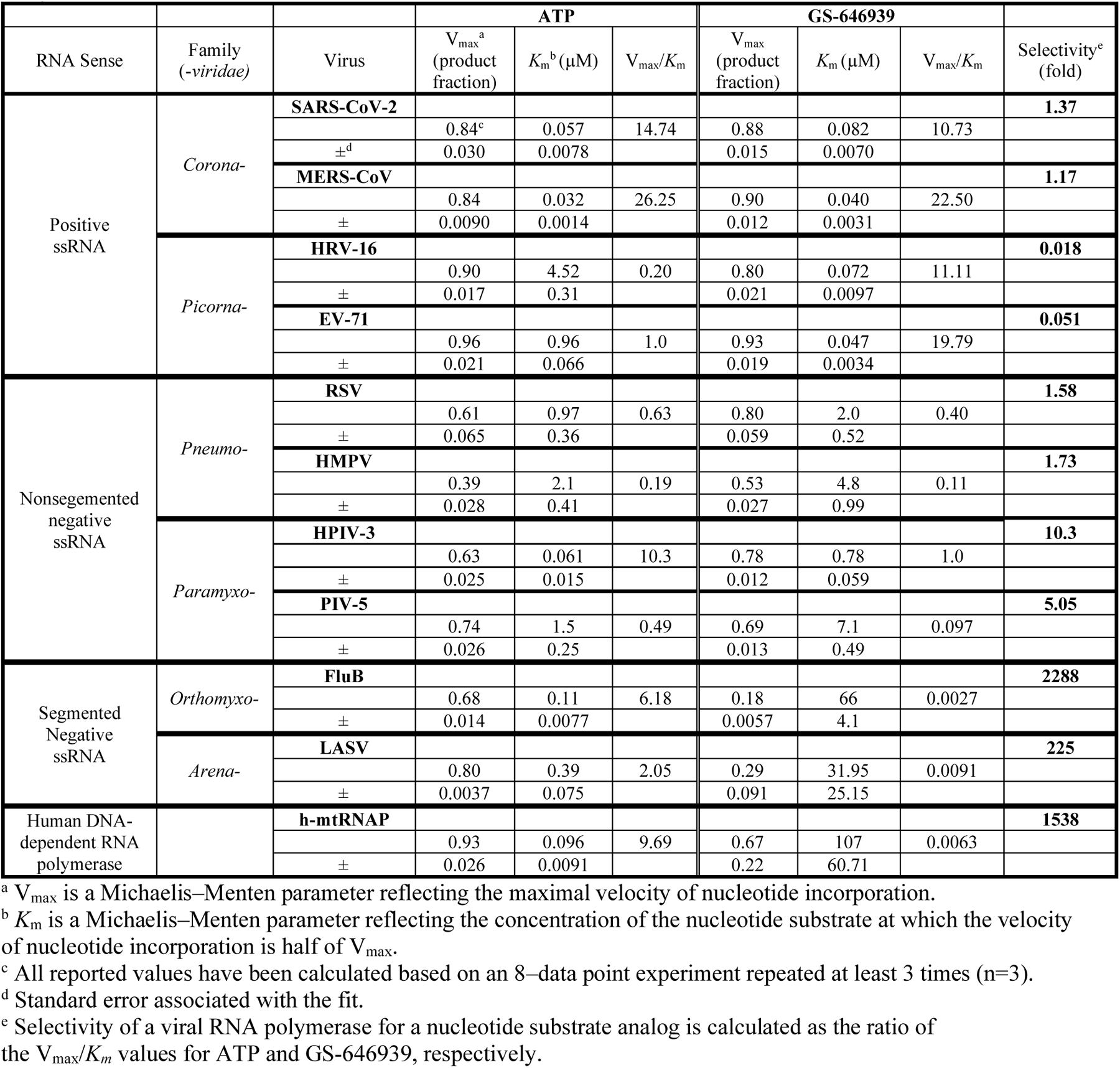
Selective incorporation of GS-646939, a 4ʹ-cyano purine NTP analog, by selected RdRp enzymes.

GS-646939 shows incorporation selectivity values of approximately 1.2-1.4 for SARS-CoV-2 and MERS-CoV RdRp (Table 3). For HRV-16 and EV-71 RdRp, GS-646939 selectivity values are as low as 0.018, markedly exceeding the incorporation efficiency of the 1ʹ-cyano GS-443902 by up to ∼50-fold. RSV and HMPV RdRp show selectivity values of ∼1.5 for GS-646939, a slight improvement over GS-443902. Incorporation by HPIV-3 and PIV-5 RdRp show comparable selectivity values between 5 and 10 for both nucleotide analogs. Similar to GS-443902, GS-646939 served as a poor substrate for influenza B and LASV RdRp, generating selectivity values greater than 200. Likewise, h-mtRNAP demonstrated poor incorporation efficiency of GS-646939 with a selectivity value of ∼1500. Overall, GS-646939 and GS-443902 follow similar trends. The exception is the preference for GS-646939 by picornavirus RdRp and a predilection for GS-443902 by coronavirus RdRp.

The exceptional substrate utilization of GS-646939 by HRV-16 and EV-71 RdRp suggests that the nucleotide analog can outcompete its natural counterpart ATP. To address this question directly, we simultaneously added ATP and GS-646939 to the reaction and monitored incorporation of the corresponding monophosphate (MP) formed (Fig. 2). AMP (“i_1_”) and GS-646939 (“i_2_”) terminated primers could be distinguished from one another due to the difference in migration pattern (Fig. 2*A*). The concentration of GS-646939 required to match 50% ATP incorporation is defined as the matching concentration (MC_50_). As expected for a competitive inhibitor, the MC_50_ value increased with increasing concentrations of the competing ATP (Fig. 2*B*). The ratio of the MC_50_ value over the ATP concentration provides the competitive index (CI) (Fig. 2*C*). The lower the value, the better the competitive advantage for incorporation of a given nucleotide. The average CI values for GS-646939 were 0.04 and 0.07 for HRV-16 and EV-71 RdRp, respectively (Fig. 2*D*). These data suggest that, under competitive conditions, the picornavirus RdRp enzymes use GS-646939 ∼14-to 25-fold more efficiently than ATP. Furthermore, both picornavirus enzymes demonstrated a greater than 30-fold preference for GS-646939 over GS-443902 under competitive conditions (Fig. S2). Together, the competitive incorporation data corroborate the selectivity measurements.

**Figure 2:**
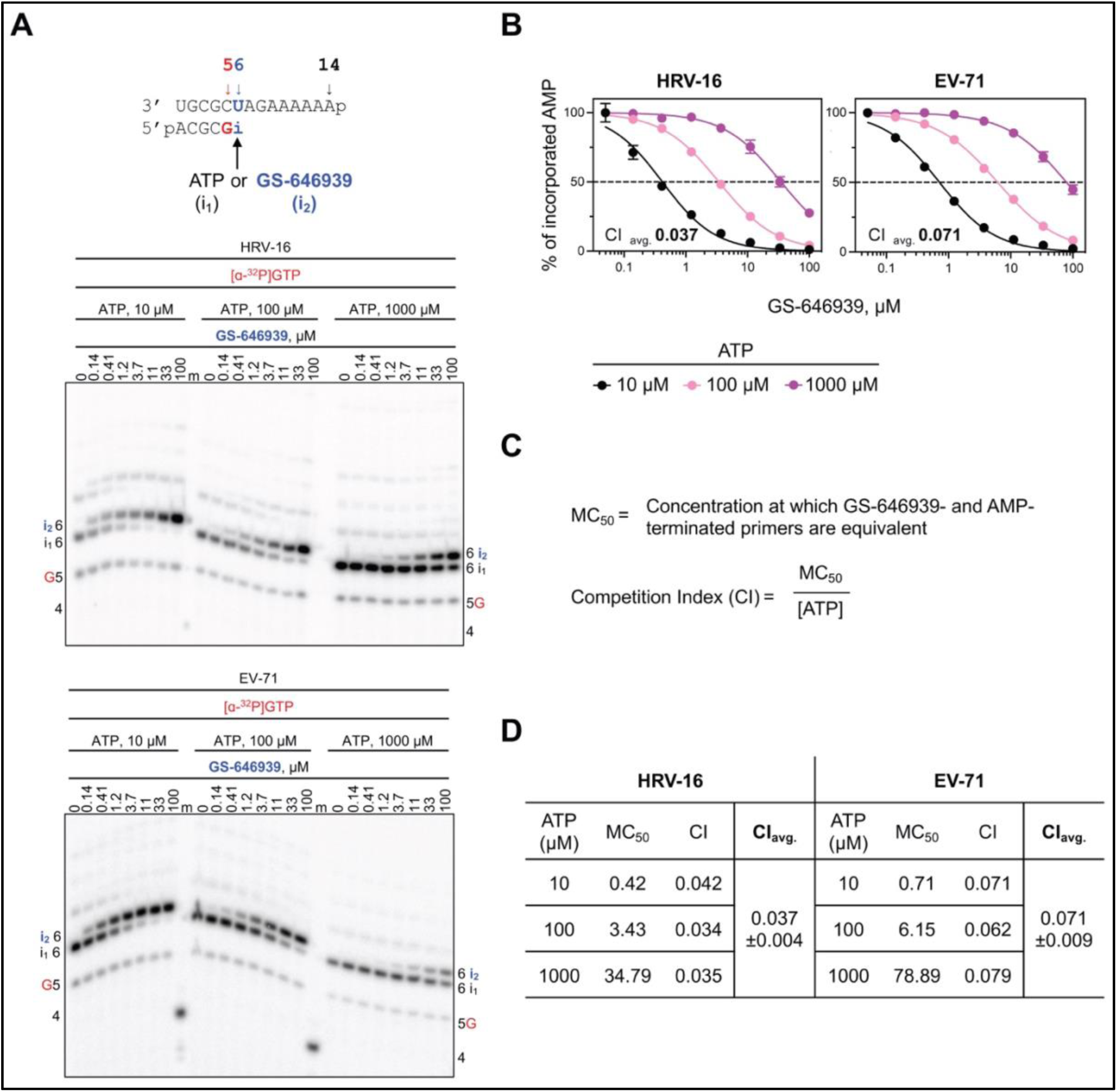
Incorporation of GS-646939 under competitive conditions by HRV-16 and EV-71 RdRp. ***A***, RNA primer/template (*top*) supporting a single incorporation of ATP (“i_1_”) or GS-646939 (“i_2_”) at position 6. G indicates incorporation of [α-^32^P]-GTP at position 5. Migration pattern of RNA synthesis products catalyzed by HRV-16 (*middle*) and EV-71 RdRp (*bottom*). Product formation resulting in AMP-or GS-646939-terminated primers was compared across increasing GS-646939 concentrations at ATP concentrations of 10, 100, and 1000 µM. A 5′-^32^P-labeled 4-nt primer serves as a size marker. ***B***, Graphical representation of AMP-terminated primers (%) at increasing GS-646939 concentrations as shown in ***A***. Independent 8-data point experiments were performed at least three times (n=3) and error bars represent standard error associated with the fit. ***C***, The MC_50_ value is defined as the concentration at which GS-646939 matches ATP for incorporation at position 6. The competition index (CI) is the ratio of the MC_50_ value to the ATP concentration present in the reaction. ***D***, CI values determined for HRV-16 and EV-71 RdRp and the CI average (CI_avg_) and standard deviation (±) across all ATP concentrations.

### Structural models of incorporation

Based on existing x-ray and cryo-EM structures (25,36,38,40–43), we generated models of the pre-incorporated states of ATP, GS-443902, and GS-646939 for representative RdRp enzymes investigated in this study (Table 1). The active site of viral RdRps is generally characterized by a set of motifs with highly conserved residues involved in substrate binding and catalysis (48). However, subtle variations in the active site landscape seem to govern the specificity of inhibitor incorporation. For HRV-16, EV-71, SARS-CoV-2, MERS-CoV, RSV, HMPV, PIV-5 and HPIV-3, both the 1ʹ-cyano of GS-443902 and the 4ʹ-cyano of GS-646939 are tolerated to varying degrees. As previously described, the 1ʹ-cyano of GS-443902 is particularly well positioned in the coronavirus active site, occupying a uniquely polar pocket defined by Thr-687, Asn-691 and Ser-759 in SARS-CoV-2 (25), and similarly for MERS-CoV. For GS-646939, the 4′-cyano appears to have a favorable interaction with Asn-296 of HRV-16 (Fig. 3*A*). The Motif C residue Gly-326 in HRV-16 allows for maximum flexibility in the 4ʹ pocket, in contrast to the similar SARS-CoV-2 and MERS-CoV pocket, in which the corresponding residue is a serine (Fig. 3*B*). For RSV (Fig. 3*C*) and HMPV, PIV-5 (Fig. 3*D*) and HPIV-3, the 4ʹ-cyano fits nicely in the pocket, but doesn’t appear to convey any advantage.

**Figure 3.**
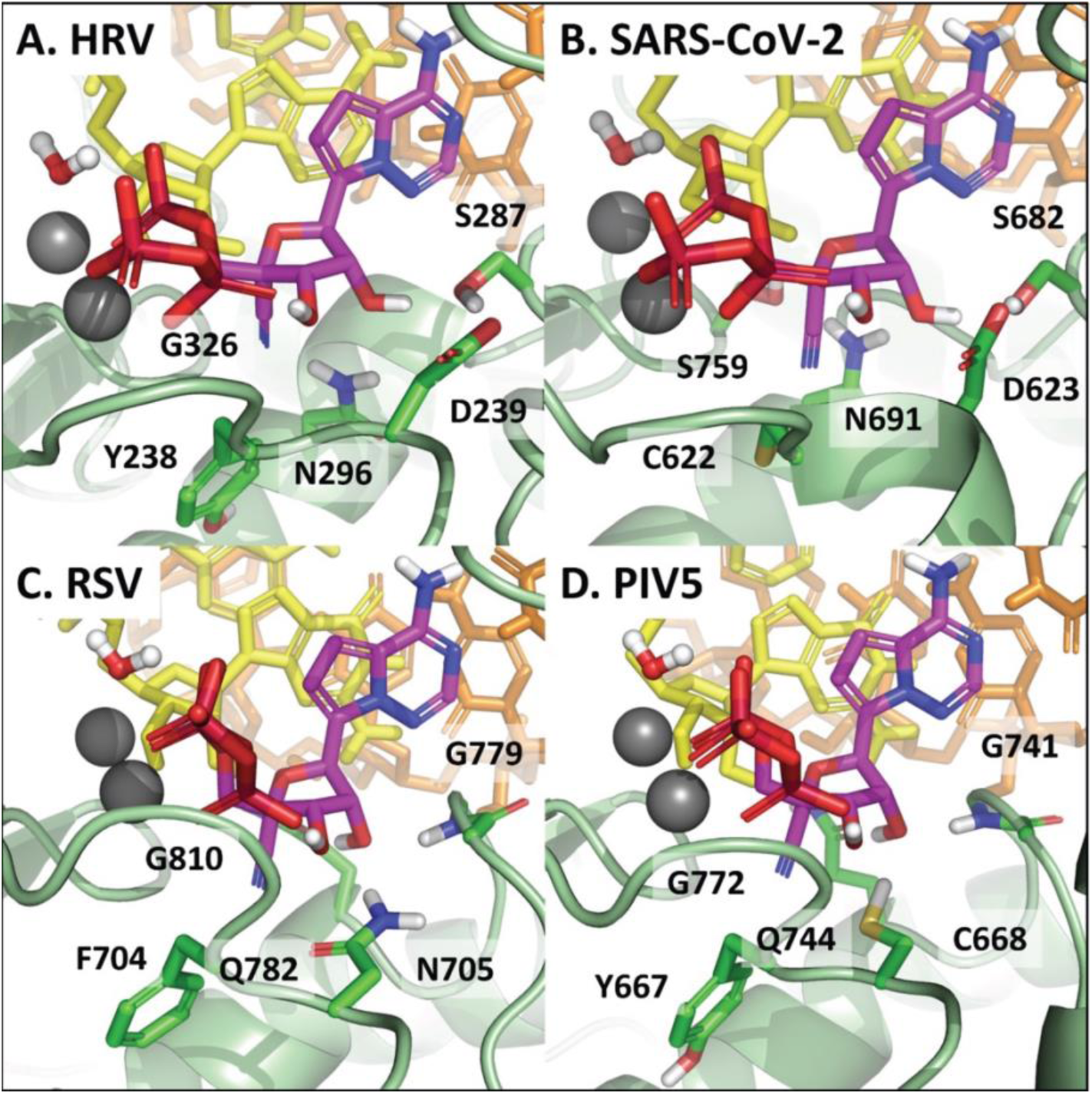
Models of GS-646939 in its pre-incorporated state for ***A***, HRV, ***B***, SARS-CoV-2, ***C***, RSV, and ***D***, PIV5. Selectivity is largely driven by the nature of the 4ʹ pocket and the specifics of how the 2ʹ-OH is recognized by the polymerase. HRV and SARS-CoV-2 have a similar overall active site structure, differing primarily at Gly-326/Ser-759 and Tyr-238/Cys-622. While somewhat different from HRV and SARS-CoV-2, RSV and PIV5 also have a similar overall structure, differing primarily at Phe-704/Tyr-667 and Asn-705/Cys-668. In each case, the 4ʹ-cyano of GS-646939 is at least tolerated. The interaction between the 4ʹ-cyano and Asn-296 in HRV appears to be particularly ideal and is likely responsible for the inhibitor’s increased affinity compared to ATP.

For LASV and FluB, both the 1ʹ-cyano of GS-443902 and 4ʹ-cyano of GS-646939 disrupt a water-mediated hydrogen bond network, which is essential for recognition of the substrate 2ʹ-OH (Fig. S3*A* and Fig. S3*B*). For GS-646939, the water molecule is likely completely displaced, with no means to compensate for the loss in hydrogen bonding at 2′. Compounding the issue, both LASV and FluB have a bulky tryptophan residue forming the floor of the 4ʹ pocket. The comparable residue in the polymerases for which GS-646939 has demonstrably greater activity is typically a tyrosine, phenylalanine, or another smaller residue. This provides a degree more flexibility to accommodate the 4ʹ-cyano. In the case of h-mtRNAP, Motif B is fundamentally different from viral RdRps (Fig. S3*C*). With respect to GS-443902, the 1ʹ pocket is clearly occluded by His-1125, while for GS-646939, the 4ʹ pocket is occluded by a salt bridge formed from Arg-802 and Asp-1128. In both cases, incorporation of the inhibitor should be significantly compromised.

### Inhibition of RNA primer extension reactions

Incorporation of the nucleotide analog into the growing RNA chain is a prerequisite for any potential inhibitory effect. Notably, GS-443902 and GS-646939 possess a 3ʹ-hydroxyl group, classifying them as non-obligate chain-terminators. The presence of a 3ʹ-hydroxyl group may allow for the nucleophilic attack on the incoming nucleotide and its subsequent incorporation. Previous biochemical and structural studies of SARS-CoV-2 have established that incorporation of GS-443902 at position “i” results in delayed-chain termination at position “i+3” (26–28). This mechanism can be attributed to a steric clash between the 1ʹ-cyano group and the hydroxyl group of the conserved Ser-861. A structural comparison between Nsp12 of SARS-CoV-2 and the 3D^pol^ of HRV-16 and EV-71 revealed that the picornaviruses share an analogous serine (Fig. 4*A*) (29). RNA synthesis following GS-443902 incorporation by SARS-CoV-2, EV-71, and HRV-16 RdRp at position “i” generated the same intermediate product at position “i+3” (Fig. 4*B*). As reported for SARS-CoV-2 RdRp, the inhibitory effect was favoured at low nucleotide concentrations and could be overcome with increasing concentrations of the nucleotide substrate at position “i+4” (26,27).

**Figure 4:**
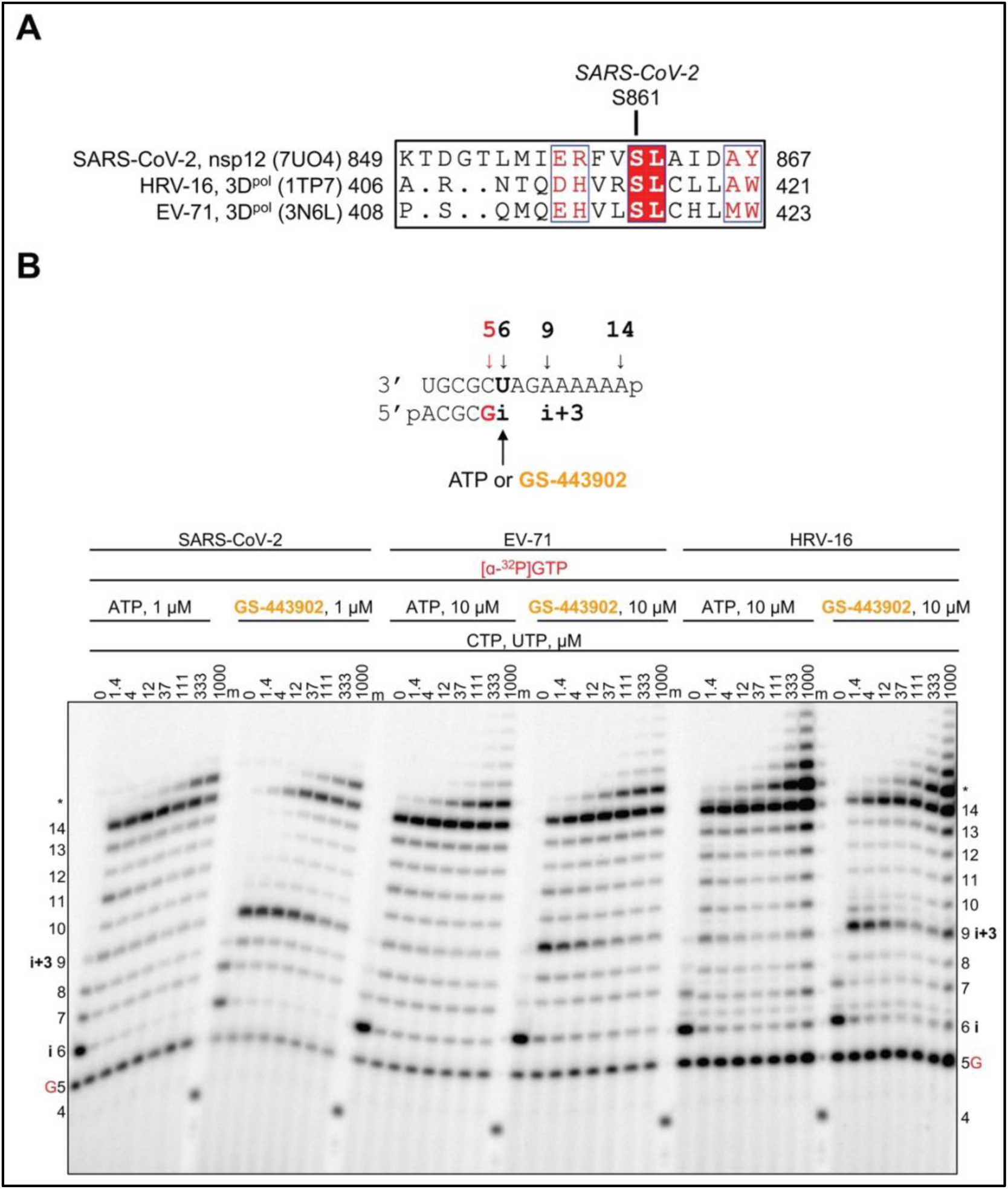
SARS-CoV-2, EV-71, and HRV-16 RdRp-catalyzed RNA synthesis pattern of inhibition following a single incorporation of ATP or GS-443902 as a function of nucleotide concentration. *A*, Sequence alignment based on a 3D structural overlay of SARS-CoV-2 Nsp12 (PDB:7UO4), HRV-16 3D^pol^ (PDB:1TP7), and EV-71 3D^pol^ (PDB:3N6L) composed using ESPript 3.0 (80). Ser-861 (SARS-CoV-2 numbering) is conserved in HRV-16 and EV-71 RdRp. *B*, RNA primer/template supporting a single incorporation of ATP or GS-443902 at position 6 (*top*). G indicates incorporation of [α-^32^P]-GTP at position 5. Extension following the incorporation of ATP and GS-443902 at position 6 (“i”) at increasing CTP and UTP concentrations catalyzed by SARS-CoV-2, EV-71, and HRV-16 RdRp (*bottom*). Following GS-443902, an intermediate product forms at position 9 (“i+3”), which is overcome at elevated CTP and UTP concentrations. A 5′-^32^P-labeled 4-nt primer serves as a size marker. Product formation at and above the *asterisk* indicates RNA products that are likely a result of sequence-dependent slippage events.

Employing the same approach as above, we show that GS-646939 inhibited HRV-16 and EV-71 RdRp-mediated RNA synthesis at the site of incorporation (Fig. 5). While GS-443902 incorporation promotes delayed chain-termination, GS-646939 causes immediate chain-termination. Inhibition by GS-646939 is overcome with increasing NTP concentration following the incorporated analog. A similar pattern was observed for SARS-CoV-2 and MERS-CoV RdRp (Fig. S4). GS-443902 inhibits other RdRp enzymes likewise via delayed chain-termination. However, the patterns are more heterogeneous, with variations in the position of inhibition; the structural reasons have yet to be determined (46). In contrast, GS-646939 incorporation by RSV and HMPV RdRp resulted in immediate chain-termination (Fig. S5), which was nearly absolute and could not be overcome to a discernable extent with increased NTP concentrations. Inhibition of HPIV-3 and PIV-5 RdRp with GS-646939 shows a similar immediate chain-termination pattern (Fig. S6).

**Figure 5:**
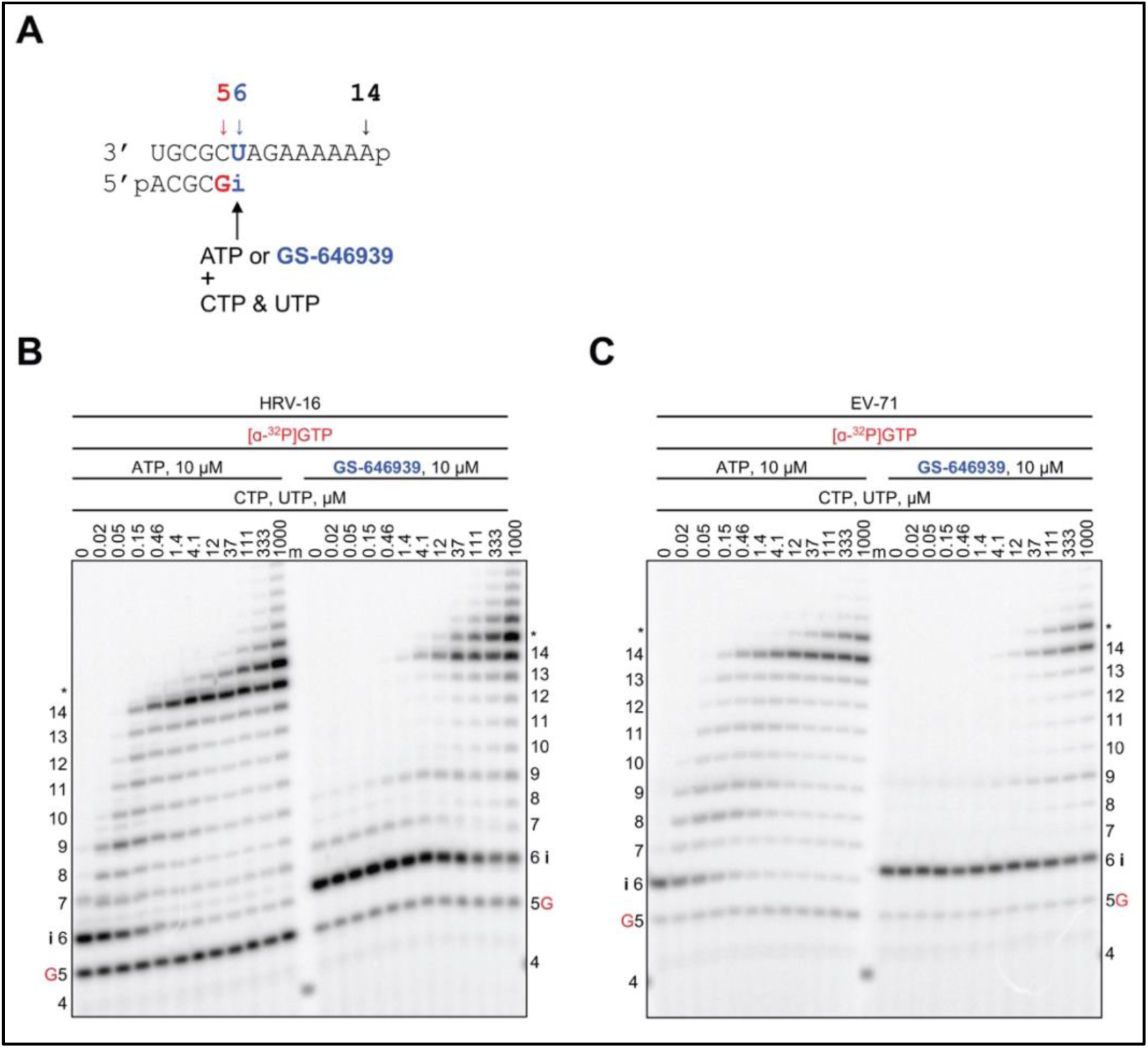
HRV-16 or EV-71 RdRp-catalyzed RNA synthesis and pattern of inhibition following a single incorporation of ATP or GS-646939 as a function of nucleotide concentration. *A*, RNA primer/template supporting RNA synthesis and a single incorporation of ATP or GS-646939 at position 6 (“i”). G indicates incorporation of [α-^32^P]-GTP at position 5. *B*, Migration pattern of RNA products resulting from HRV-16 RdRp-catalyzed RNA extension of AMP (*left*) of GS-646939 (*right*) at increasing concentrations of CTP and UTP. A 5′-^32^P-labeled 4-nt primer serves as a size marker. Product formation at and beyond the asterisk indicates RNA products that are likely a result of sequence-dependent slippage events. *C*, RNA synthesis products catalyzed by EV-71 RdRp. Incomplete inhibition of RNA synthesis occurs at the site of GS-646939 incorporation (“i”), full template-length product is generated at elevated nucleotide concentrations.

To quantify the inhibitory effect of GS-443902 and GS-646939 on subsequent nucleotide incorporation, we generated AMP-, GS-443902-, and GS-646939-terminated primers and measured kinetic parameters of the natural UTP substrate incorporation at position “i+1” (Fig. 6 and Table S1). For all RdRp enzymes, compared to AMP-terminated primers, the extension of GS-646939-terminated primers required a marked increase in UTP concentration. Conversely, GS-443902-terminated primers promoted subtle inhibition of UTP incorporation for HRV-16, EV-71, RSV, and HMPV RdRp. For SARS-CoV-2, MERS-CoV, HPIV-3, and PIV-5 RdRp, equivalent or improved utilization of the GS-443902-terminated primer was observed.

**Figure 6:**
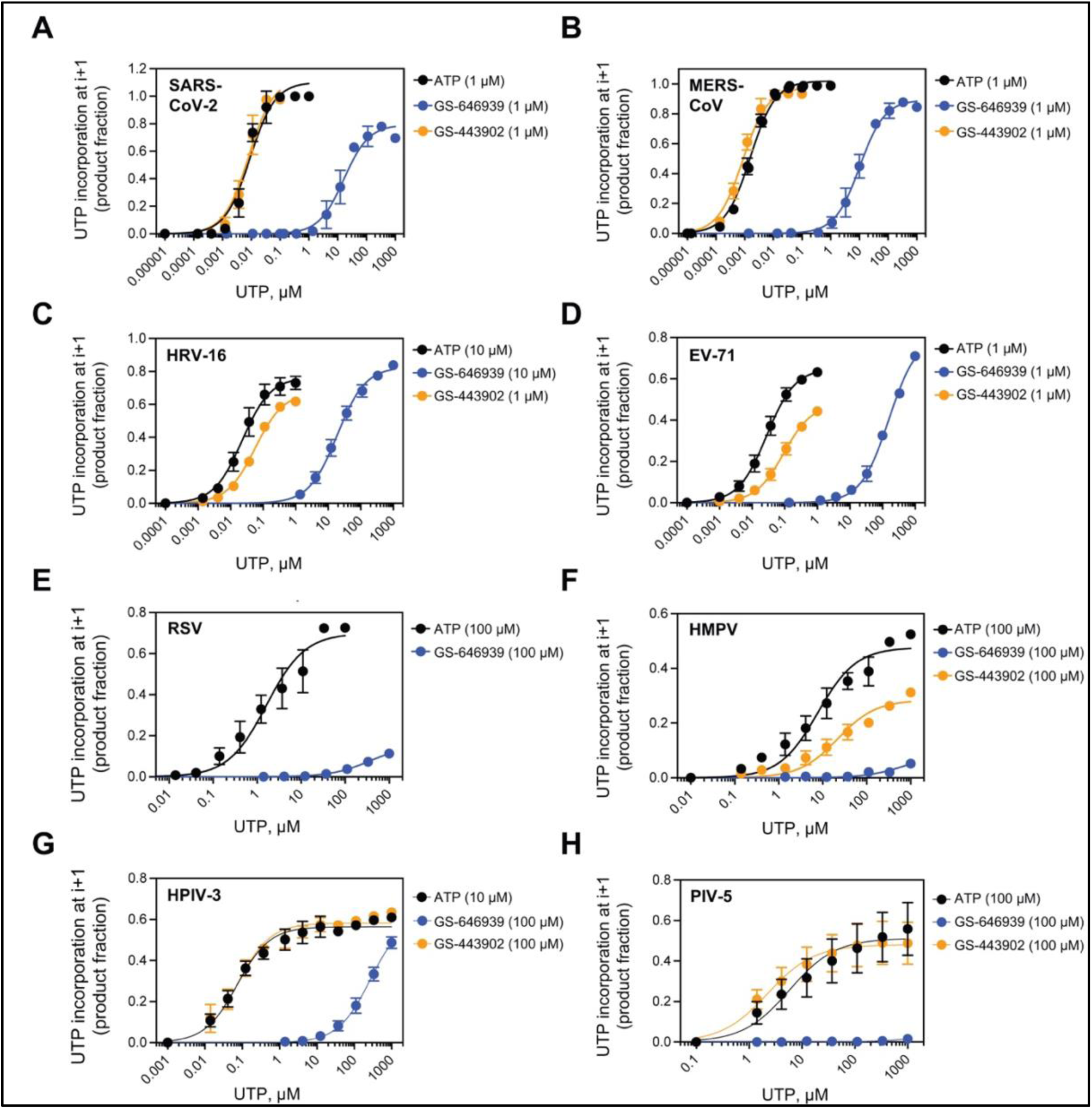
GS-646939 inhibits subsequent nucleotide incorporation. UTP incorporation at position “i+1” was monitored at increasing concentrations immediately following ATP (*black*), GS-443902 (*green*), or GS-646939 (*blue*) incorporation at position “i”. Concentrations of ATP, GS-443902, and GS-646939 supplemented to the reaction are shown in brackets. Viral RdRp enzymes investigated include SARS-CoV-2 (***A***), MERS-CoV (***B***), HRV-16 (***C***), EV-71 (***D***), RSV (***E***), HPIV-3 (***G***), and PIV-5 (***H***). The product fraction was calculated as the total signal above position “i” divided by total signal in the lane. Independent 8-data point experiments were performed 3 times (n=3) and error bars represent standard error associated with the fit.

### Structural rationale for chain-termination

An analysis of the trajectory of the incorporated GS-646939 during translocation from the “i” substrate position to the “i+1” primer position may help to provide a better understanding of the requirements for chain-termination. For all the RdRps studied here, the 4ʹ-cyano moiety of the inhibitor must pass through a set of Motif C residues, which coordinate the two catalytic metals (Fig. 7). The obstacle presented by these residues is largely independent of the specific sidechains and instead derives from the protein backbone itself.

**Figure 7.**
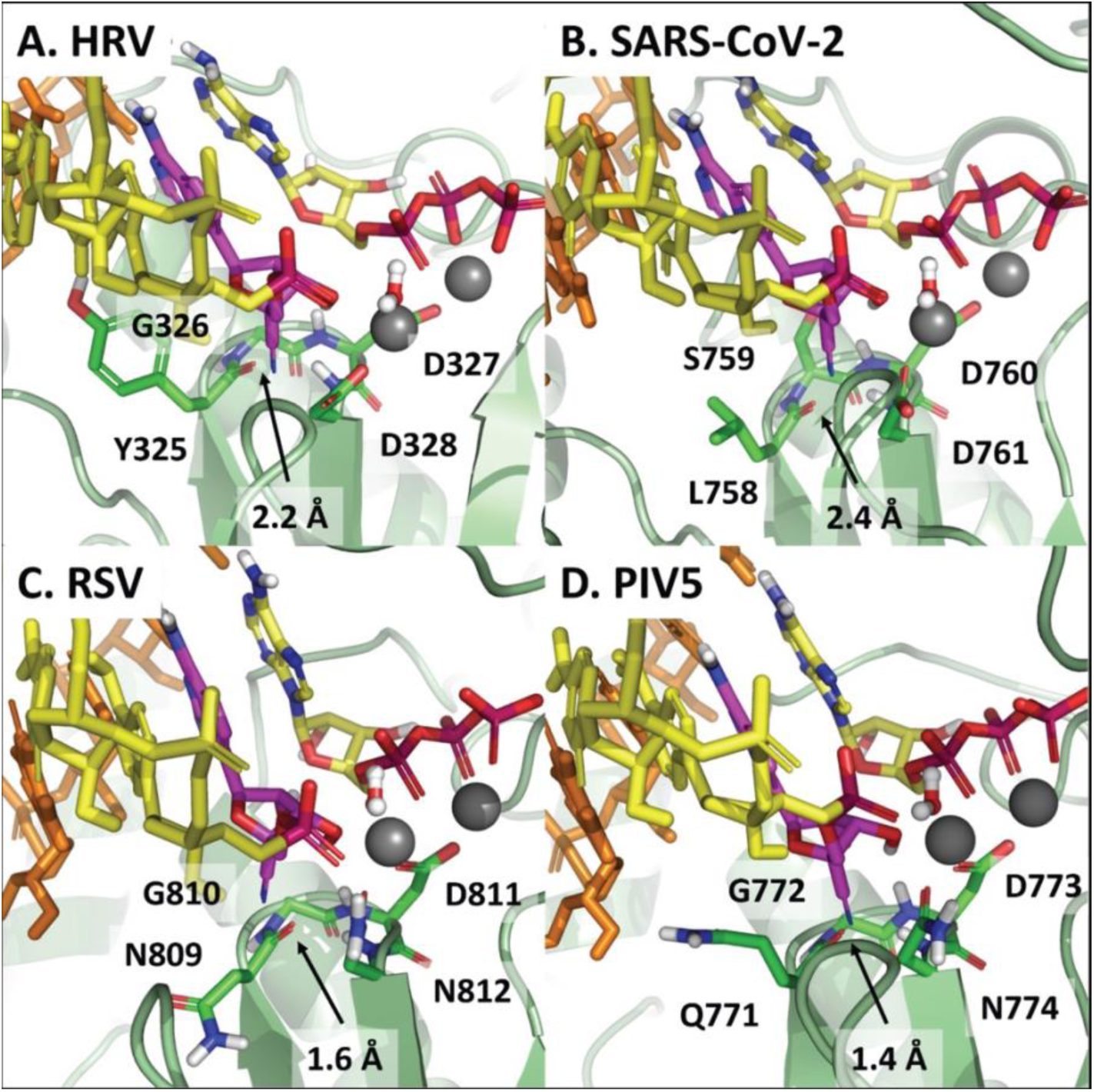
Models of incorporated GS-646939 translocated to the “i+1” position in ***A***, HRV, ***B***, SARS-CoV-2, ***C***, RSV, and ***D***, PIV5. Structures are not optimized but serve as a guide to clashes that impair further incorporation. Following incorporation, translocation of the RNA to position the inhibitor in the first primer position is hindered by residues of Motif C which coordinate the catalytic metals. For HRV, Cα of Gly-326 and NH of Asp-327 present a translocation obstacle to the 4ʹ-cyano of the incorporated inhibitor. A similar impediment to translocation exists for the other RdRps. Assuming translocation occurs, C=O of Tyr-325 presents a significant steric clash which would perturb proper primer positioning. This clash appears to be more significant in RSV and PIV5.

For HRV (Fig. 7*A*) and SARS-CoV-2 (Fig. 7*B*), this clash is moderate. But for RSV (Fig. 7*C*) and PIV5 (Fig. 7*D*), the clash appears to be severe, significantly impairing proper positioning of the primer. If translocation is indeed compromised, one would expect diminished NTP incorporation independent of the nature of the incoming nucleotide. Like GS-646939, 4ʹ-ethynyl-2-fluoro-2ʹ-deoxyadenosine (EFdA) possesses a bulky 4ʹ-modification and was shown to block translocation of human immunodeficiency virus type 1 reverse transcriptase (HIV-1 RT) (49–51). Therefore, we used EFdA as a benchmark. For HRV-16 RdRp, GS-646939-and EFdA-terminated primers generally reduce NTP incorporation of all nucleotides tested (Fig. S7). High concentrations of UTP and 2-thio-UTP are required to obtain the “i+1” product, while 3ʹdeoxy UTP, 2ʹO-methyl UTP, and 2’deoxy UTP are not incorporated. In contrast, AMP-and GS-443902-terminated primers do not cause immediate chain-termination and instead enable the incorporation of each of these nucleotide analogs to a similar extent.

### Significance of template-dependent inhibition by GS-646939

For SARS-CoV-2 RdRp, it has been demonstrated that higher NTP concentrations can overcome GS-443902-induced delayed chain-termination or pausing (23,26,27,31). This in turn may lead to full-length products containing embedded nucleotide analogs. When used as a template, UTP incorporation opposite the complementary GS-443902 is diminished. This unified template-dependent inhibition mechanism for GS-443902 has been described for several viral RdRps (27). Here, we designed RNA templates to monitor UTP incorporation opposite a template-embedded GS-443902 or GS-646939 residue, respectively (Fig. 8*A*). For the HRV-16 RdRp, submicromolar UTP concentration was sufficient to generate full-template length product on Template “A” that contains the natural nucleotide (Fig. 8*B*). Comparatively, on Template “GS-443902” and “GS-646939”, inhibition opposite the embedded analog could be rescued by increasing UTP concentrations ∼8- and 2-fold, respectively (Fig. 8*C*). Similarly, EV-71 RdRp-catalyzed UTP incorporation was inhibited to a greater extent across GS-443902 (∼16-fold) than GS-646939 (∼9-fold) (Fig. S8). For SARS-CoV-2 and MERS-CoV RdRp, RNA synthesis opposite GS-646939 generates multiple intermediate products, indicative of an inhibitory effect (Fig. S9). To conclude, GS-443902 and GS-646939 cause inhibition in primer extension reactions when embedded in the template. While the template-dependent mechanism seems to be the dominant mode of inhibition of RdRp enzymes by GS-443902, GS-646939 acts primarily as a chain-terminator.

**Figure 8:**
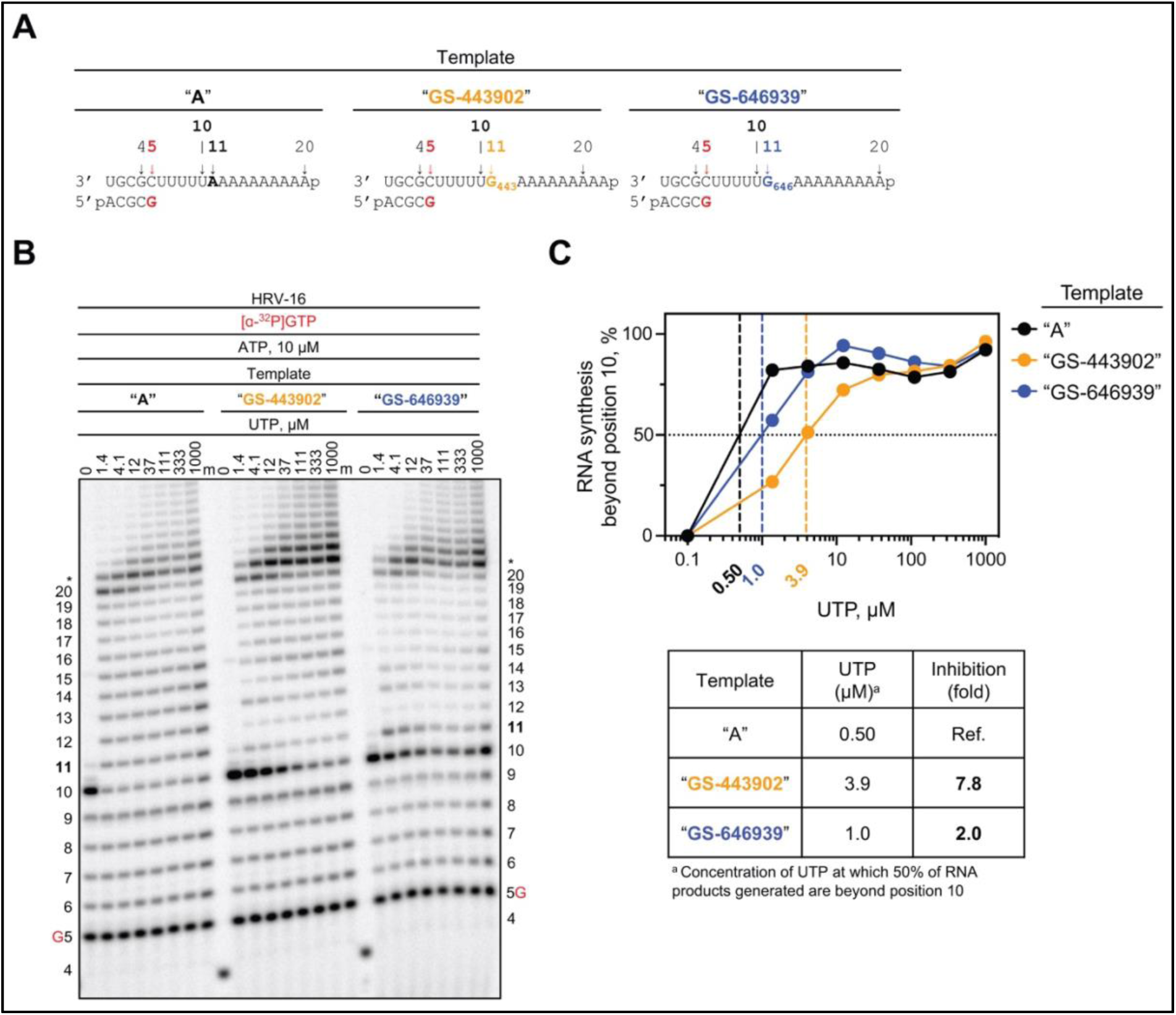
RNA synthesis catalyzed by HRV-16 RdRp using a template with a single GS-443902 or GS-646939 residue embedded in the template at position 11. *A*, RNA primer/template with an embedded GS-443902 (*Template “GS-443902”, middle*) or GS-646939 (*Template “GS-646939”, right*); the corresponding primer/template with adenosine (*Template A*) at this position is shown on the left. G5 indicates incorporation of [α-^32^P]-GTP at position 5. *B*, Migration pattern of the products of RNA synthesis catalyzed by HRV-16 RdRp. MgCl_2_, [α-^32^P]-GTP, and ATP provided to the reaction to support RNA synthesis up to position 10. Increasing concentrations of UTP were supplemented to the reactions to monitor incorporation opposite a templated adenosine, GS-443902, or GS-646939 at position 11, and templated adenosines from position 12 to 20. Compared to Template A, intermediate products form at position 10 on Template “GS-443902” and “GS-646939”, indicating template-dependent inhibition. Product formation at and beyond the asterisk indicates RNA products that are likely a result of sequence-dependent slippage events. A 5′-^32^P-labeled 4-nt primer serves as a size marker. *C*, Quantification of *B* (*top*) where the sum of RNA products generated beyond position 10 was divided by the total signal in the lane and normalized as a percentage, fold-inhibition resulting from an embedded GS-443902 or GS-646939 (*bottom*). To account for template-dependent differences in activity, product fraction was normalized as a percentage to product fraction observed at 1000 µM UTP for that template.

## DISCUSSION

The availability of effective antivirals for treating infections with respiratory RNA viruses is limited. Nucleotide analog RdRp inhibitors have the potential to act broadly against diverse RNA viruses, which is of particular importance in outbreak situations with emerging pathogens. Mutagenic nucleotides such as ribavirin, favipiravir, or molnupiravir provide prominent examples in this regard (52–57). These compounds are base-modified, which can cause lethal mutagenesis when the active triphosphate metabolite is used as a substrate (45,58). Classic nucleotide analogs with modifications in the sugar moiety commonly inhibit RNA synthesis. Here, we studied the mechanism of action of the GS-646939 nucleotide triphosphate metabolite of the newly discovered GS-7682 4ʹ-cyano modified *C*-adenosine nucleotide phosphoramidate analog. Inhibition of RNA synthesis was evaluated against an array of purified recombinant RdRp enzymes representing medically relevant respiratory RNA viruses. The broadly acting 1ʹ-cyano modified *C*-adenosine nucleotide triphosphate analog GS-443902 was included in this study for comparative purpose. Despite a certain degree of overlap in the spectrum of antiviral activities, the results of this study demonstrate distinct mechanisms of action for GS-443902 and GS-646939. While GS-443902 inhibits RNA synthesis predominantly when embedded in the template strand, GS-646939 causes chain-termination at the site of incorporation.

For RNA polymerases evaluated in this and our previous studies, the data reveal a decrease in selectivity for GS-443902 in the order of SARS-CoV-2, SARS-CoV, MERS-CoV > HRV-16, EV-71 > RSV, HMPV > HPIV-3, PIV-5 > LASV > influenza B ≫ h-mtRNAP. GS-443902 is a better substrate for the SARS-CoV-2 RdRp complex than its natural counterpart ATP (24,26). Cryo-EM structures of the enzyme complex with bound RNA and a pre-incorporated GS-443902 revealed a hydrophilic pocket composed of Nsp12 Thr-687, Asn-691, and Ser-759 that accommodates the 1ʹ-cyano group (25). Of note, the S759A mutation was shown to confer resistance to GS-5734, due to a 5-to 10-fold increase in discrimination against GS-443902 (25,47). Thus, biochemical and structural data align with *in vitro* selection experiments in explaining the selectivity for GS-443902 over ATP.

The preference for selective incorporation of GS-646939 follows the order: HRV-16 and EV-71 > SARS-CoV-2 and MERS-CoV ≈ RSV and HMPV > HPIV-3 and PIV-5 ≫ LASV ≫ influenza B = h-mtRNAP. For HRV-16 and EV-71 RdRp, GS-646939 is used ∼20-50 fold more efficiently as a substrate than ATP. To the best of our knowledge, this is the most favourable selectivity ever reported for the incorporation of a nucleotide analog by viral polymerases. Modeling of the pre-incorporated GS-646939 indicates that the 4ʹ-cyano is tolerated by each of the aforementioned polymerases with the exception of LASV, influenza B and h-mtRNAP. With respect to HRV-16 and EV-71 RdRp, the particular combination of residues that form both the 2ʹ-OH recognition motif and the 4ʹ pocket allow for a favorable interaction between the 4ʹ-cyano and Asn-296 in HRV. In the case of LASV and influenza B, both the 1ʹ-cyano of GS-443902 and the 4ʹ-cyano of GS-646939 disrupt a water-mediated hydrogen bonding network which serves to recognize the ribose 2ʹ-OH. The active site of h-mtRNAP is dissimilar to any of the viral RdRps, with predicted clashes for both 1ʹ-cyano and 4ʹ-cyano substitutions.

GS-443902-mediated inhibition of primer extension reactions is generally heterogeneous (44). The position and extent of inhibition depend upon the nature of the polymerase, and the underlying structural determinants have not yet been investigated for RSV and other RdRps. In contrast, all viral RNA polymerases tested towards this end inhibit the incorporation of UTP opposite an embedded GS-443902 in the template. This inhibition may not necessarily translate into significant antiviral effects because the incorporation of GS-443902 may be inefficient in some cases, as it is for LASV and influenza viruses (44). For GS-646939, the unifying mechanism of action is chain-termination. The extent of inhibition depends on the NTP concentration that can overcome blockages. Chain-termination with HRV-16 and EV-71 RdRp is indeed not absolute, and higher NTP concentrations allow the continuation of RNA synthesis. However, the high selectivity for GS-646939 over ATP seems sufficient to cause potent antiviral effects. Chain-termination with SARS-CoV-2 and MERS-CoV is also overcome with higher NTP concentrations, but a limited antiviral effect in cellular assays may be explained by the presence of a viral exonuclease, which can remove the incorporated inhibitor and allow synthesis to continue (59). For RSV and HMPV, chain-termination is nearly absolute and lower nucleotide analog incorporation rates are therefore still sufficient for potent antiviral effects.

The antiviral effects and biochemical properties of several other nucleotide analogs with bulky 4ʹ-modifications have been reported, and chain-termination is the dominant mechanism of action (51,60–67). Prodrugs of 4ʹ-azidocytidine (balapiravir) and 4ʹ-chloromethyl-2ʹ-deoxy-2ʹ-fluoro-cytidine (lumicitabine) were developed for the treatment of HCV and RSV infection, respectively. EFdA was developed for the treatment of HIV-1 infection. EFdA-TP is efficiently incorporated by HIV-1 RT, and structural data provide strong evidence to show that chain-termination is based on compromised enzyme translocation (49–51). A hydrophobic pocket accommodates the 4ʹ-ethynyl moiety, which facilitates binding of EFdA-TP and stabilizes the complex after incorporating the nucleotide analog. The favourable interactions in the pre-translocated state and unfavourable interactions in the post-translocated state essentially block RT translocation and further incorporation events. Our structural models of RdRp enzymes considered in this study point to a similar mechanism with GS-646939. In contrast, 4’-fluorouridine was shown to cause delayed chain-termination against RSV RdRp (60). This data suggests that the relatively small 4’-fluoro modification does not interfere with translocation at the point of incorporation.

In conclusion, 1ʹ-cyano and 4ʹ-cyano modified nucleotide analogs inhibit diverse RdRp enzymes of respiratory RNA viruses via different mechanisms. Enzymes of the *Coronaviridae* (SARS-CoV-2 and MERS-CoV) and the *Pneumoviridae* (RSV and HMPV), are effectively targeted by the 1ʹ-cyano modified *C*-adenosine GS-443902. Selective incorporation and efficient template-dependent inhibition provide biochemical explanations for the observed antiviral activity (23,26,27). RdRps of the *Picornaviridae* (HRV-16 and EV-71) and *Pneumoviridae* (RSV and HMPV) are effectively targeted by the 4ʹ-cyano modified *C*-adenosine GS-646939. Selective incorporation and efficient chain-termination were identified here as important parameters that correlate with potent antiviral activity. Taken together, the reported data provide strong support for the prototypic pathogen approach at the biochemical level. Here we studied the effects of antiviral agents against the polymerase of two prototypic pathogens within a given family of RNA viruses. The results provide confidence that the 1ʹ-cyano and 4ʹ-cyano modified nucleotides can be considered broadly to target emerging viruses that belong to these families.

## MATERIALS AND METHODS

### Nucleic acids and chemicals

RNA primers and templates used in this study were 5′-phosphorylated and purchased from Dharmacon (Lafayette, CO, USA). GS-646939 and GS-443902 were provided by Gilead Sciences (Foster City, CA, USA). NTPs were purchased from GE Healthcare (Mississauga, ON, Canada). [α-^32^P]GTP was purchased from PerkinElmer (Revvity, Waltham, MA,USA). 3′deoxy UTP, 2′O-methyl UTP, and 2-thio UTP were purchased from TriLink Biotechnologies (San Diego, CA, USA). 2′deoxy UTP was purchased from MedChem Express (Monmouth Junction, NJ, USA).

### Expression and purification of viral polymerases

Expression and purification of MERS-CoV, SARS-CoV-2, RSV, LASV, FluB, and h-mtRNAP used in this study have been described (23,26,27). The baculovirus expression system was employed for HMPV, HPIV-3, and PIV-5 RdRp P/L complex. The pFastBac-1 (Invitrogen, Burlington, ON, Canada) plasmid with the codon-optimized synthetic DNA sequences (GenScript, Piscataway, NJ, USA) coding for HMPV (L: AAQ67700 and P: AAQ67693), were used as starting material for protein expression in insect cells (Sf9, Invitrogen, Burlington, ON, Canada). HMPV P/L complex was expressed as a polyprotein in frame with the N-terminal Tobacco etch virus (TEV) protease using the MultiBac (Geneva Biotech; Geneva, Switzerland) (68,69). This approach was originally reported for the expression of influenza virus RdRp trimeric complex (70,71). P and L proteins were cleaved post-translationally from the polyprotein at the engineered TEV sites. HPIV-3 (L: NP_067153.2 P: 067149.1), and PIV-5 (L: AFE48451.1 and P: AFE48445.1) were cloned into the pFastBac Dual vector (Invitrogen, Burlington, ON, Canada), P was under the control of the p10 promoter and L under the polyhedrin promotor. The RdRp P/L complexes were purified using Ni-NTA affinity chromatography based on their respective eight-histidine tag as shown in **Table 1** (Thermo Scientific, Rockford, IL, USA). The pET-15b (Novagen) plasmid with codon-optimized synthetic DNA sequences (GenScript, Piscataway, NJ, USA) coding for the RdRp of EV-71 (BAJ49823.1) and HRV-16 (AAA6982.1) were expressed in *Escherichia coli* BL21. EV-71 and HRV-16 RdRp enzymes were purified using Ni-NTA affinity chromatography based on their respective C-terminal eight-histidine tag according to the manufacturer’s specifications (Thermo Scientific). The protein identities of the purified HMPV, HPIV-3, and PIV-5 RdRp P/L complex as well as EV-71 and HRV-16 RdRp were confirmed by mass spectrometry analysis (Alberta Proteomics and Mass Spectrometry Facility, University of Alberta, Canada).

### Evaluation of GS-646939 and GS-443902 incorporation and subsequent primer- and template-strand inhibition on viral RNA synthesis

The following synthetic 5′-monophosphorylated RNA templates were used in this study (the portion of the template that is complimentary to the 4-nt primer is underlined): 3′<u>TGCG</u>CTAGTTT for h-mtRNAP selectivity values; 3′<u>UGCG</u>CUAGAAAAAAp for MERS-CoV, SARS-CoV-2, EV-71, HRV-16, FluB, LASV analog selectivity, ATP/GS-646939 and GS-646939/GS-443902 competition, pattern of inhibition, and UTP and UTP analog incorporation at position “i+1” experiments; 3′<u>UGCG</u>CUAGUUUAUUp for RSV, HMPV, HPIV-3, and PIV-5 analog selectivity, pattern of inhibition, and UTP incorporation at position “i+1” experiments; 3′<u>UGCG</u>CUUUUUAAAAAAAAAAp, 3′<u>UGCG</u>CUUUUUG_646_AAAAAAAAAp, and 3′<u>UGCG</u>CUUUUUG_443_AAAAAAAAAp were used for the evaluation of nucleotide incorporation opposite GS-646939 (Template “GS-646939”) and GS-443902 (Template “GS-443902”). RNA synthesis assays of viral RdRp, data acquisition, and quantification were performed as previously reported by us (23,26,27,44,45,47,72). Briefly, viral RdRp concentration was optimized to ensure incorporation of [α-^32^P]-GTP is within the linear range. For the evaluation of single nucleotide incorporation, the concentration range of ATP, GS-646939, and GS-443902 were optimized to avoid misincorporation at subsequent positions. Reaction mixture for RNA synthesis assay (final concentrations after mixing), contain the purified viral RdRp, Tris–HCl (pH 8, 25 mM), RNA primer (200 μM), RNA template (2 μM, except for FluB RdRp reactions where optimal RNA template concentration was 0.5 µM), [α-^32^P]-GTP (0.1 µM), and various nucleotide concentrations were prepared on ice and incubated at 30℃ for 5 to 10 minutes. RNA synthesis was initiated by the addition of 5 mM MgCl_2_, except for SARS-CoV-2 and MERS-CoV RdRp, which required 1.25 mM MgCl_2_. The duration of reactions varied amongst the enzymes, SARS-CoV-2, MERS-CoV, HRV-16, EV-71, and h-mtRNAP reactions were stopped after 10 minutes; for RSV, HMPV, HPIV-3, PIV-5, FluB, and LASV reactions were stopped after 30 minutes. All reactions were stopped by the addition of equal volume formamide/EDTA (50mM). The reaction products were incubated at 95℃ for 5 minutes and then resolved by 20% UREA-PAGE, and the [α-^32^P] generated signal was stored and scanned using Typhoon phosphorimager (Cytiva, Vancouver, BC, Canada).

The data were analyzed using GraphPad Prism 7.0 (GraphPad Software, Inc; San Diego, CA, USA).

### Generation of structural models

Models of nucleotide incorporation for representative enzymes in this study were generated based on existing x-ray or cryo-EM structures. For SARS-CoV-2, models were based on the cryo-EM structure 7UO4 (25), which includes a template, a 3’deoxy-primer, pre-incorporated GS-443902, and a single Mn^++^ ion. A 3’OH was added to the primer, the Mn^++^ was changed to Mg^++^ and a second metal was added. The active site was then refined with Macromodel (Schrödinger, LLC, New York, USA) (73,74) for incoming ATP, GS-646939 and GS-443902. The HRV-A16 model was based on the x-ray structure 4K50 (36), which includes only primer and template. Based on the SARS-CoV-2 model and an x-ray structure for the similar HCV polymerase (4WTD) (75), the two metals were added, and the structure was refined for the same three NTP’s as above. The Lassa model was based on the cryo-EM structure 7OJN (42), which includes template, primer, two Mn^++^ ions, and a non-hydrolyzable UTP analog. In this case, some ambiguity in how the NTP 2’OH was recognized by the enzyme suggested to us that a bridging water may be involved. To answer this question, a WaterMap (Schrödinger, LLC, New York, USA) (76,77) analysis was run, which indicated that a single water molecule linked the NTP 2’OH to the active site residues His-1298 and Ser-1330. A similar situation arose for FluB, which was based on the x-ray structures 6QCV and 6QCX (41). In this case, we found a single water molecule linked the NTP 2’OH to the active site residues Asn-414 and Ser-443. The model for h-mtRNAP was based on the x-ray structure 4BOC (43), which includes primer and template. Again, we took a similar approach to optimizing for incoming ATP, GS-443902 and GS-646939 as above. More challenging were the models for RSV (based on the apo cryo-EM structure 6PZK (38) and PIV5 (based on the apo cryo-EM structure 6V85 (40)). These structures required major modifications to accommodate the primer, template, metals and NTP. Similarity in the structure to Lassa allowed us to insert those elements from that model into the apo structure for RSV. The protein was then optimized around the RNA, using sidechain sampling and minimization in Prime (Schrödinger, LLC, New York, USA) (78,79), before allowing the RNA to also relax using Macromodel. Sidechain conformations in the active site were modified following the precedent of the above structures to recognize the metals and the NTP. Notably, active site Motif A (residues 700-705) underwent a significant conformational change from its apo form. Investigation of multiple sidechain conformations to recognize the NTP 2’OH ultimately settled on a configuration in which Met-705 and Gln-782 directly hydrogen bond to the ribose. No bridging water molecule was predicted. The optimization of the PIV5 model followed a similar path.

## Data availability

All data are included within this article

## Acknowledgements

The authors would like to thank Emma Woolner for excellent technical assistance and Dr. Jack Moore at the Alberta Proteomics and Mass Spectrometry facility for mass spectrometry analysis.

## Author contributions

CJG, SMW, EPT, DK, and JKP investigation; CJG, SMW, and EPT validation; CJG, SMW, EPT, JKP, and MG formal analysis; CJG, SMW, EPT, and JKP visualization; CJG and MG writing–original draft; CJG, SMW, EPT, DK, JP, DSS, JKP, JYF, JPB, and MG. writing–review and editing; JYF, JPB, and MG resources; CJG data curation; MG conceptualization; CJG, SMW, EPT, JKP, and MG methodology; MG supervision; MG project administration; MG funding acquisition.

## Funding and additional information

This study was supported by grants to MG from the Canadian Institutes of Health Research (CIHR) under the funding reference number 170343, Gilead Sciences Inc., the Alberta Ministry of Technology and Innovation through SPP-ARC (Striving for Pandemic Preparedness - The Alberta Research Consortium) and by the National Institute of Allergy and Infectious Diseases of the National Institutes of Health under Award Number U19AI171292. The content is solely the responsibility of the authors and does not necessarily represent the official views of the National Institutes of Health. CG is supported by a grant from the CIHR under the funding reference number 181545.

