## Supporting Information for "Mechanism and spectrum of inhibition of a 4’-cyano modified nucleotide analog against diverse RNA polymerases of prototypic respiratory RNA viruses"

Supplemental Table 1: Inhibitory effect of the incorporated analog-monophosphate on incorporation of the subsequent nucleotide

| Virus | Primer 3'-end (base) | Substrate | $V_{max}$[b] (product fraction) | $K_m$[c] (μM) | $V_{max}/K_m$ | Inhibition[d] (fold) |
|---|---|---|---|---|---|---|
| **SARS-CoV-2** | AMP | UTP | 1.10 [e] ±[f] 0.043 | 0.009 ± 0.0015 | 122 | Ref [g] |
| | GS-646939 (MP[a]) | | 0.79 ± 0.036 | 16.42 ± 2.67 | 0.048 | **2542** |
| | GS-443902 (MP) | | 1.13 ± 0.047 | 0.0082 ± 0.0013 | 138 | **0.88** |
| **MERS-CoV** | AMP | UTP | 1.03 ± 0.016 | 0.0016 ± 0.0001 | 644 | Ref. |
| | GS-646939 (MP) | | 0.91 ± 0.10 | 24.68 ± 9.19 | 0.037 | **17,405** |
| | GS-443902 (MP) | | 0.98 ± 0.018 | 0.0009 ± 0.0001 | 1089 | **0.59** |
| **HRV-16** | AMP | UTP | 0.76 ± 0.015 | 0.023 ± 0.002 | 33.04 | Ref. |
| | GS-646939 (MP) | | 0.83 ± 0.012 | 18.88 ± 1.22 | 0.044 | **751** |
| | GS-443902 (MP) | | 0.67 ± 0.015 | 0.057 ± 0.005 | 11.75 | **2.81** |
| **EV-71** | AMP | UTP | 0.65 ± 0.003 | 0.028 ± 0.0006 | 23.21 | Ref. |
| | GS-646939 (MP) | | 0.81 ± 0.017 | 157 ± 10.25 | 0.005 | **4,642** |
| | GS-443902 (MP) | | 0.48 ± 0.005 | 0.093 ± 0.003 | 5.16 | **4.50** |
| **RSV** | AMP | UTP | 0.70 ± 0.037 | 1.6 ± 0.41 | 0.44 | Ref. |
| | GS-646939 (MP) | | 0.15 ± 0.0029 | 354 ± 16 | 0.00042 | **1,048** |
| | GS-443902 (MP) | | | | | **6.27**[1] |
| **HMPV** | AMP | UTP | 0.48 ± 0.026 | 7.8 ± 2.2 | 0.062 | Ref. |
| | GS-646939 (MP) | | 0.087 ± 0.030 | 732 ± 488 | 0.00012 | **517** |
| | GS-443902 (MP) | | 0.28 ± 0.018 | 22 ± 6.2 | 0.013 | **4.77** |
| **HPIV-3** | AMP | UTP | 0.56 ± 0.0097 | 0.073 ± 0.0093 | 7.67 | Ref. |
| | GS-646939 (MP) | | 0.61 ± 0.0099 | 268 ± 11 | 0.0023 | **3,334** |
| | GS-443902 (MP) | | 0.58 ± 0.011 | 0.076 ± 0.011 | 7.63 | **1.01** |
| **PIV-5** | AMP | UTP | 0.51 ± 0.022 | 5.8 ± 1.3 | 0.088 | Ref. |
| | GS-646939 (MP) | | 0.058 ± 0.080 | 2740 ± 4899 | 0.000021 | **4,190** |
| | GS-443902 (MP) | | 0.48 ± 0.0089 | 2.2 ± 0.26 | 0.22 | **0.40** |

[a] Denotes that the 3´-end of the primer is terminated by the monophosphate form of the respective nucleotide.
[b] $V_{max}$ is a Michaelis–Menten parameter reflecting the maximal velocity of nucleotide incorporation.
[c] $K_m$ is a Michaelis–Menten parameter reflecting the concentration of the nucleotide substrate at which the velocity of nucleotide incorporation is half of $V_{max}$
[d] Inhibition of subsequent nucleotide incorporation is calculated as the ratio of $V_{max}/K_m$ values for UTP determined on primers terminated with AMP, GS-646939, or GS-443902.
[e] All reported values have been calculated based on an 8–data point experiment repeated at least 3 times (n=3).
[f] Standard error associated with the fit.
[g] Reference.
[1] Data from Tchesnokov et al. (46)

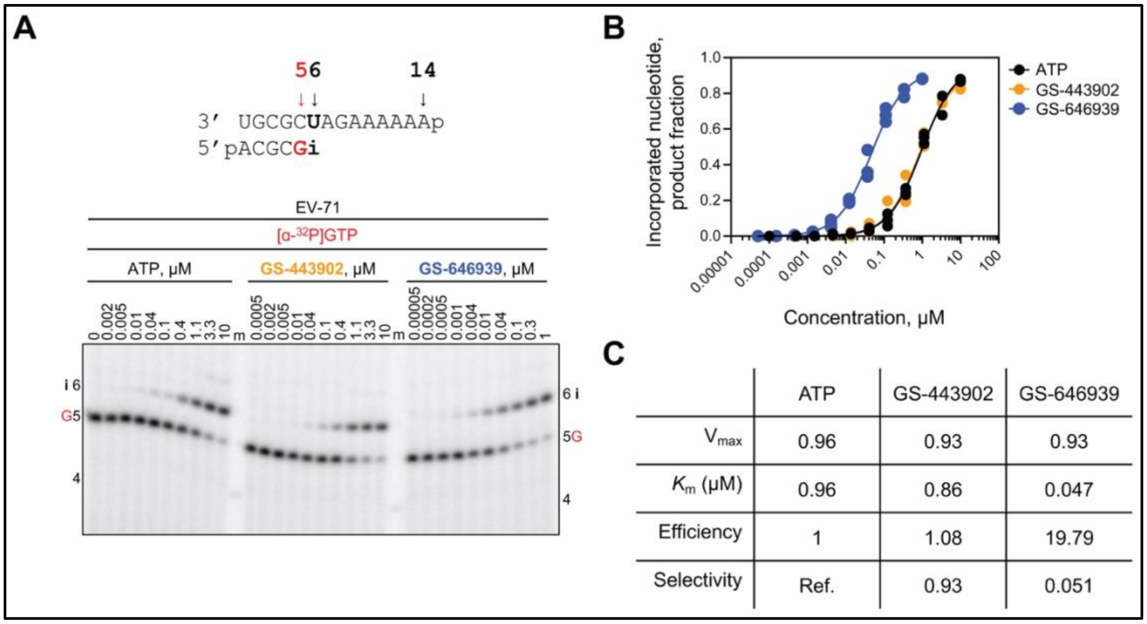


**Figure S1. Selective incorporation of GS-443902 and GS-646939 by EV-71 RdRp**. ***A***, RNA primer/template substrate used in the RNA synthesis assays to test GS-443902 and GS-646939 as substrates for incorporation at position 6 ("i") is shown above the gel. G indicates incorporation of the radiolabelled nucleotide opposite template position 5. NTP incorporation was monitored with purified EV-71 RdRp in the presence of [α-$^{32}$P]-GTP, RNA primer/template, $MgCl_2$ and increasing concentrations of ATP or the respective ATP analog substrate. Lane m illustrates the migration pattern of the radiolabelled 4 nucleotide-long primer. ***B***, Graphical representation of the incorporation of ATP and ATP analogs shown in ***A***, fitting the data points to the Michaelis-Menten function to determine $V_{max}$ and $K_m$. Independent 8-data point experiments were performed at least three times (n=3), each replicate is plotted to represent the variation between experiments. ***C***, Kinetic parameters and efficiency of ATP, GS-443902, and GS-646939 incorporation by EV-71 RdRp. Efficiency is determined as the quotient of $V_{max}$ over $K_m$ and selectivity as the quotient of natural ATP efficiency over the efficiency of the ATP analog.

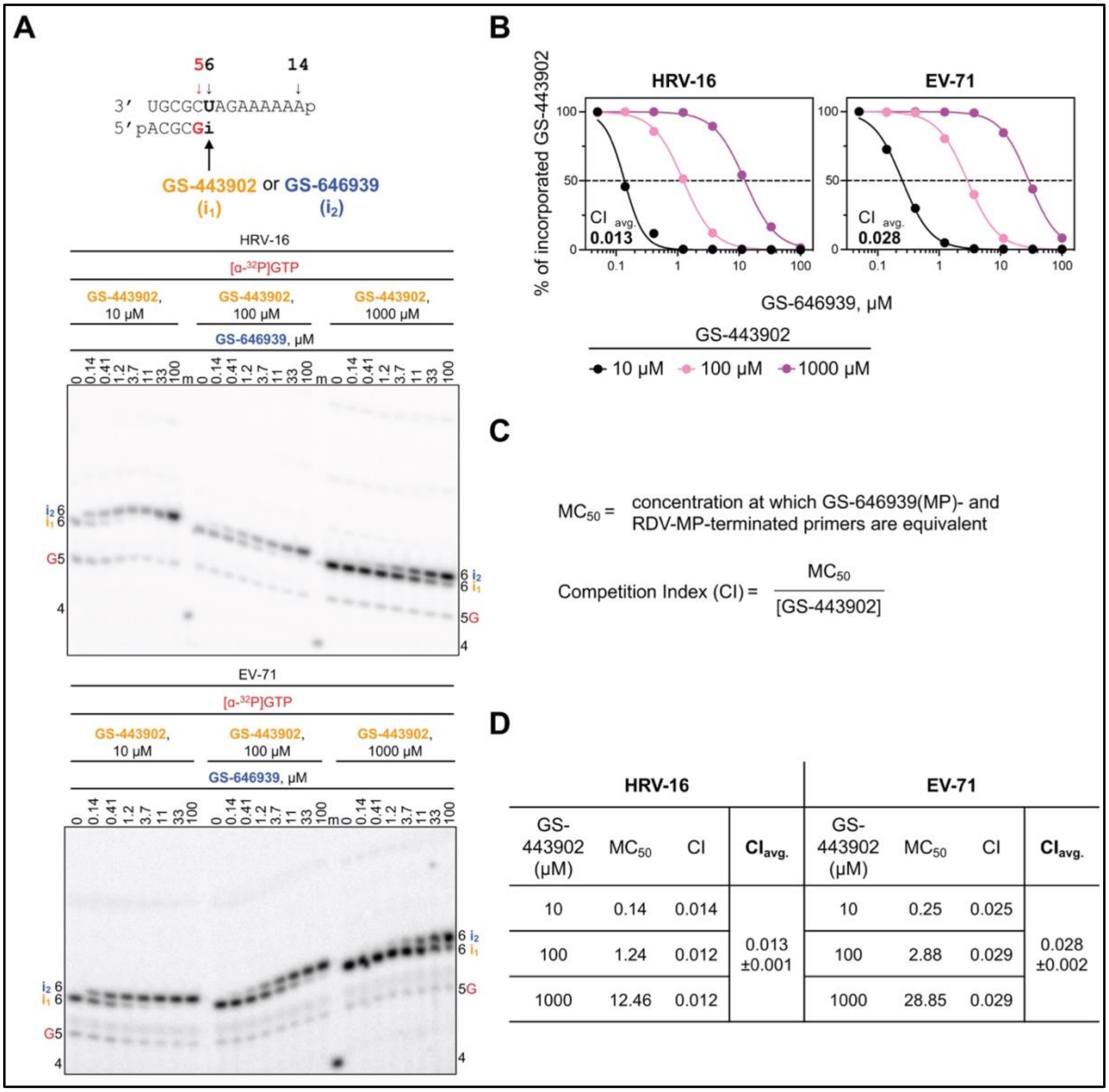


| HRV-16 | | | | EV-71 | | | |
|---|---|---|---|---|---|---|---|
| GS-443902 (µM) | $MC_{50}$ | CI | $CI_{avg.}$ | GS-443902 (µM) | $MC_{50}$ | CI | $CI_{avg.}$ |
| 10 | 0.14 | 0.014 | 0.013 ±0.001 | 10 | 0.25 | 0.025 | 0.028 ±0.002 |
| 100 | 1.24 | 0.012 | | 100 | 2.88 | 0.029 | |
| 1000 | 12.46 | 0.012 | | 1000 | 28.85 | 0.029 | |

**Figure S2: Competitive incorporation of GS-646939 versus GS-443902 by HRV-16 and EV-71 RdRp**. ***A***, RNA primer/template (top) supporting a single incorporation of GS-443902 (“$i_1$”) or GS-646939 (“$i_2$”) at position 6. G indicates incorporation of [α-$^{32}$P]-GTP at position 5. Migration pattern of RNA synthesis products catalyzed by HRV-16 (*middle*) and EV-71 RdRp (*bottom*). Product formation resulting in GS-443902- or GS-646939-terminated primers were compared across increasing GS-646939 concentrations at GS-443902 concentrations of 10, 100, and 1000 µM. A 5′-$^{32}$P-labeled 4-nt primer serves as a size marker. ***B***, Graphical representation of GS-443902-terminated primers (%) at increasing GS-646939 concentrations as shown in ***A***. ***C***, The $MC_{50}$ value is defined as the concentration at which GS-646939 outcompetes GS-443902 for incorporation at position 6. The competition index (CI) is the ratio of the $MC_{50}$ value to the GS-443902 concentration present in the reaction. ***D***, CI values determined for HRV-16 and EV-71 RdRp and the CI average ($CI_{avg}$) and standard deviation (±) across all GS-443902 concentrations.

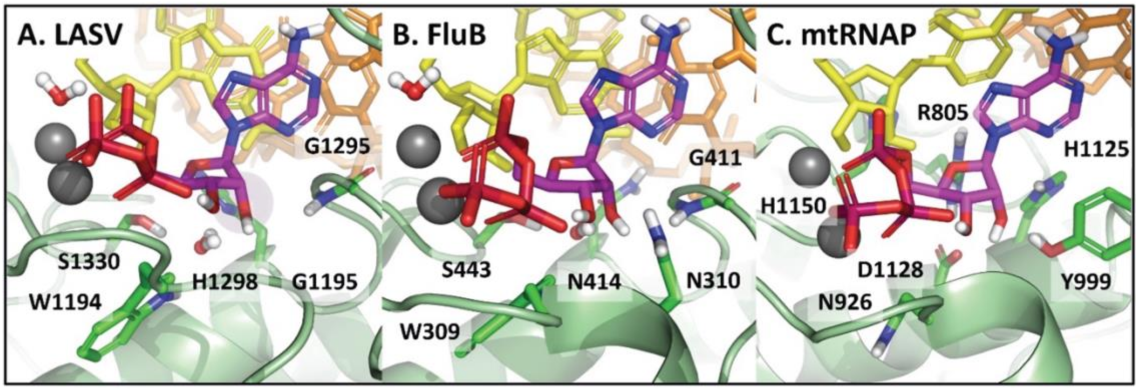


**Figure S3.** Models of ATP in its pre-incorporated state for ***A***, LASV, ***B***, FluB, and ***C***, h-mtRNAP. The active sites of LASV and FluB are similar to RSV and PIV5, but differ significantly in the specific residues which recognize the 2′-OH and define the 4’ pocket. In particular, a bridging water molecule is predicted to be a key element in the recognition of the 2′-OH. This is likely displaced by the 4′-cyano of GS-646939, compromising its affinity for these polymerases. The active site for h-mtRNAP is structurally very different from the viral polymerases. The 4′-cyano of GS-646939 appears to conflict with a salt bridge formed by Arg-805 and Asp-1128, which sit just below the ribose, making it a poor inhibitor.

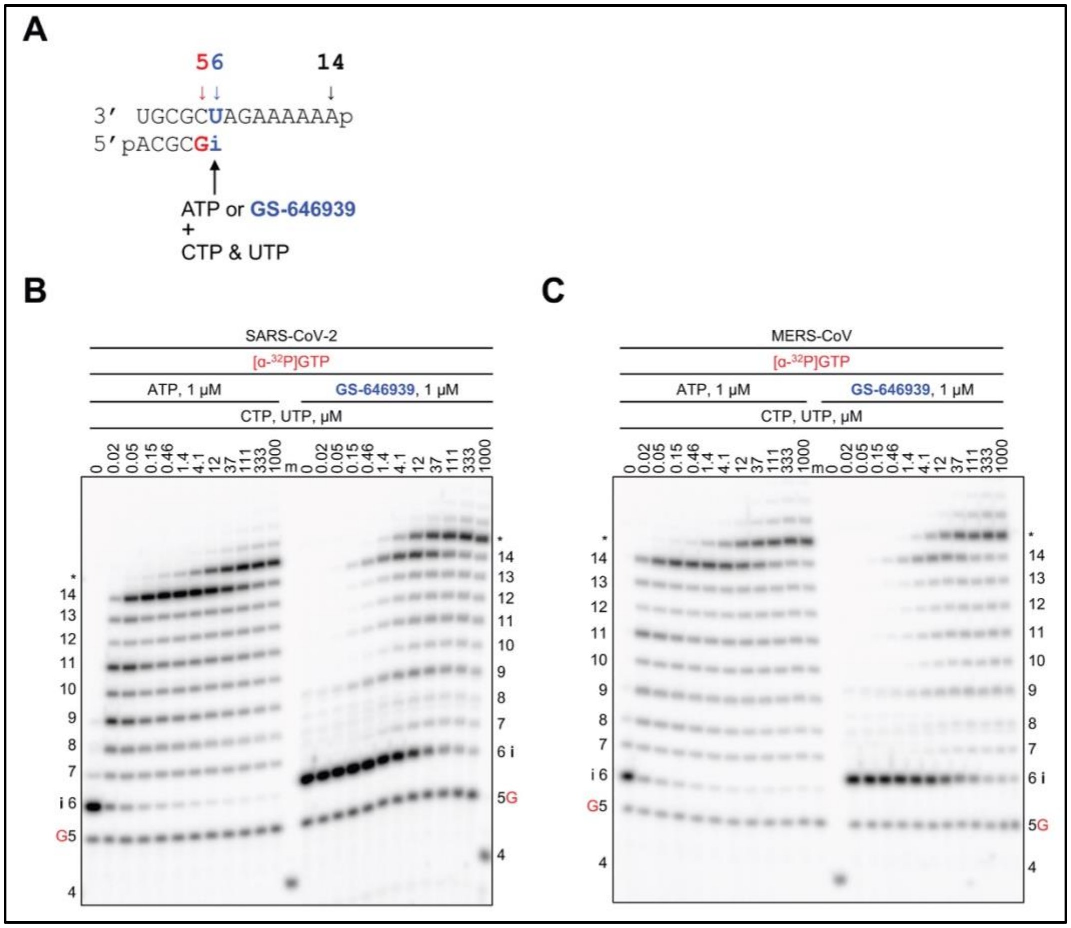


**Figure S4: SARS-CoV-2 and MERS-CoV RdRp-catalyzed RNA synthesis and inhibition patterns following a single incorporation of ATP or GS-646939 as a function of nucleotide concentration.** ***A*,** RNA primer/template supporting RNA synthesis and single incorporation of ATP or GS-646939 at position 6 ("i"). G indicates incorporation of [α-$^{32}$P]-GTP at position 5. ***B***, Migration pattern of RNA products resulting from SARS-CoV-2 RdRp catalyzed RNA extension of AMP (*left*) of GS-646939 (*right*) at increasing concentrations of CTP and UTP. A 5′-$^{32}$P-labeled 4-nt primer serves as a size marker. Product formation at and beyond the asterisk indicate RNA products that are likely a result of sequence-dependent slippage events. ***C***, Reactions with MERS-CoV RdRp, incomplete inhibition of RNA synthesis occurs at the site of GS-646939 incorporation, full template-length product is generated at elevated nucleotide concentrations.

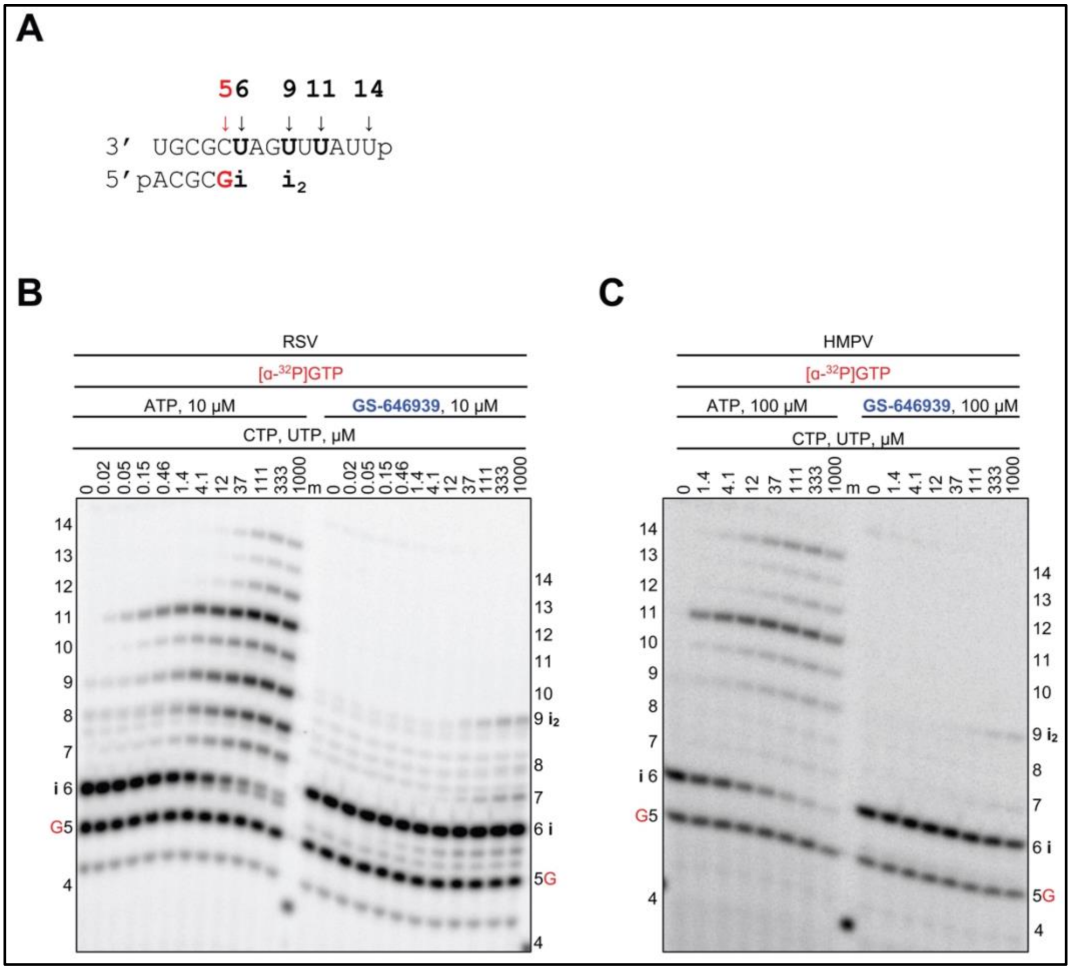


**Figure S5: RSV or HMPV RdRp-catalyzed RNA synthesis and inhibition patterns following the incorporation of GS-646939 as a function of nucleotide concentration.** ***A*****,** RNA primer/template supporting RNA synthesis and a first and second incorporation of ATP or GS-646939 by RSV and HMPV RdRp at position 6 (“i") and 9 (“$i_2$”). G indicates incorporation of [α-$^{32}$P]-GTP at position 5. ***B***, Migration pattern of RNA products resulting from RSV RdRp-catalyzed RNA extension of AMP (*left*) or GS-646939 (*right*) at increasing concentrations of CTP and UTP. A 5′-$^{32}$P-labeled 4-nt primer serves as a size marker. ***C***, Migration patterns of RNA products catalyzed HMPV RdRp, RNA synthesis was monitored at increasing CTP and UTP concentrations when ATP (*left*) or GS-646939 (*right*) were incorporated at position 6. GS-646939-terminated primers fail to be extended.

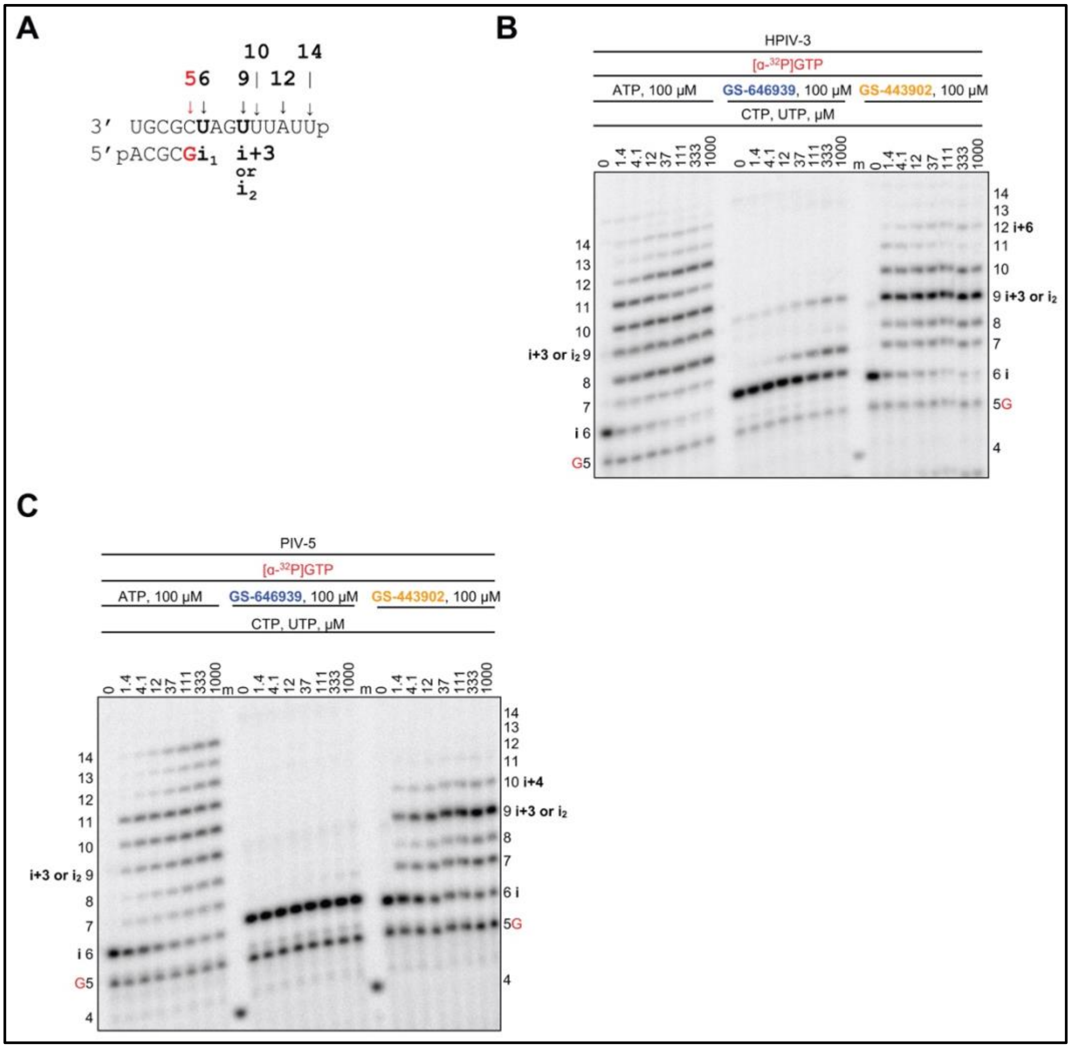


**Figure S6: HPIV-3 or PIV-5 RdRp-catalyzed RNA synthesis and inhibition patterns following the incorporation of ATP, GS-646939, or GS-443902 *A*,** RNA primer/template supporting RNA synthesis and a first and second incorporation of ATP or GS-646939 or GS-443902 at position 6 (“i”) and 9 (“$i_2$”). G indicates incorporation of [α-$^{32}$P]-GTP at position 5. ***B***, Migration patterns of RNA products catalyzed HPIV-3 RdRp. RNA synthesis was monitored at increasing CTP and UTP concentrations when natural ATP (*left*), GS-646939 (*middle*), or GS-443902 (*right*) were incorporated at position 6. GS-646939 termination was nearly absolute, whereas RNA products can be observed up to position 12 following the incorporation of GS-443902. ***C***, Migration patterns of RNA products catalyzed PIV-5 RdRp, RNA synthesis was monitored at increasing CTP and UTP concentrations when natural ATP (left), GS-646939 (*middle*), or GS-443902 (r*ight*) were incorporated at position 6. PIV-5 RdRp failed to catalyze RNA products beyond an incorporated GS-646939, while GS-443902 could be readily extended to position 9 (“i+3” or “$i_2$”).

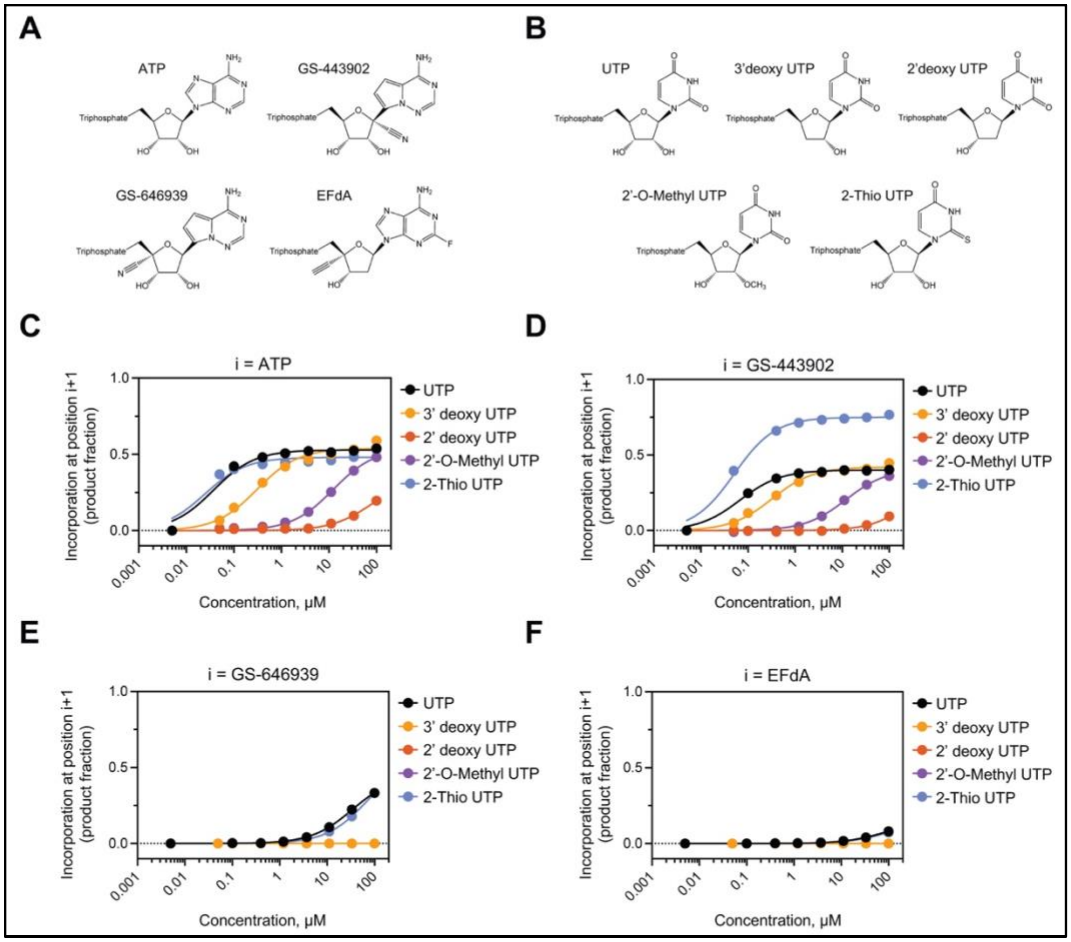


**Figure S7: GS-646939 inhibits subsequent nucleotide incorporation catalyzed by HRV-16 RdRp in a manner resembling EFdA**. ***A***, chemical structures of ATP and ATP analogs. ***B***, Chemical structures of UTP and UTP analogs. UTP and UTP analog incorporation at position 'i+1' was examined at increasing concentrations immediately following incorporation of ATP (***C***), GS-443902 (***D***), GS-646939 (***E***), or EFdA (***F***) at position "i". Product fraction was calculated as the signal generated beyond position "i" divided by the total signal in the lane.

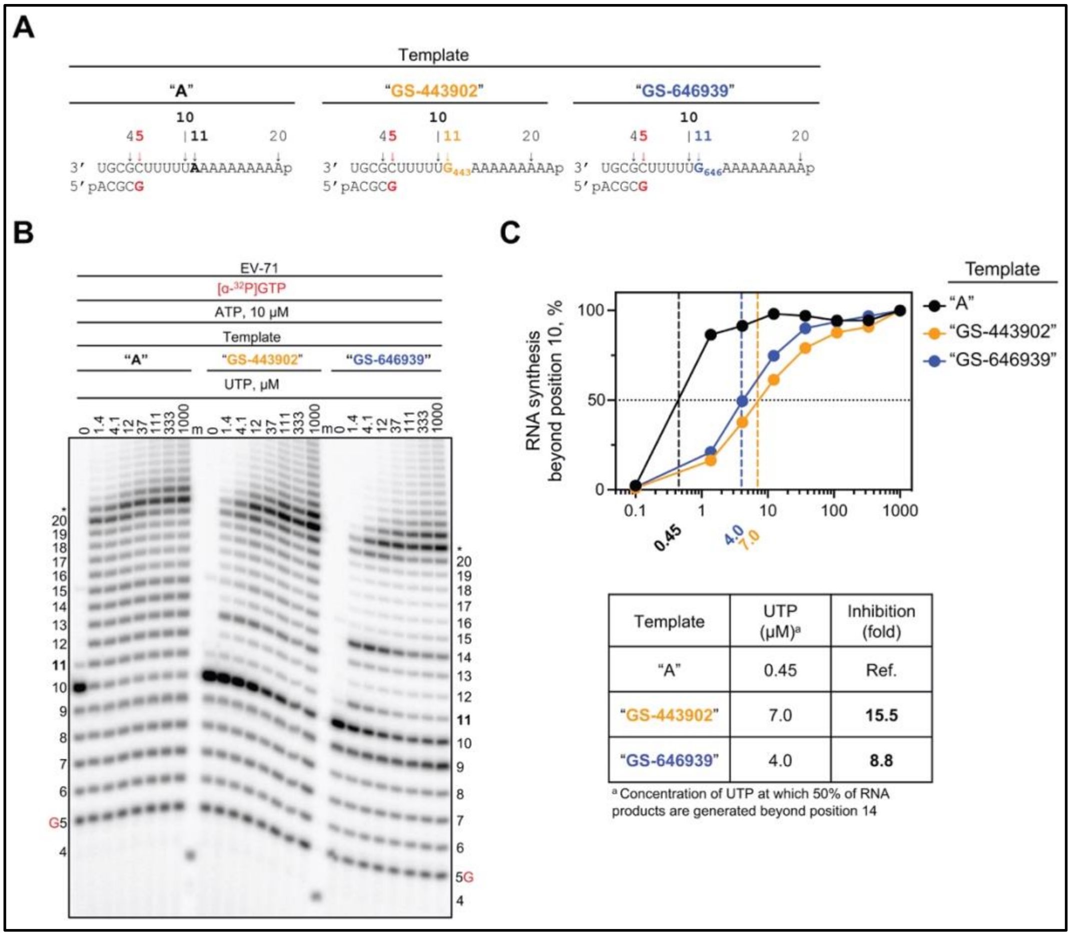


**Figure S8: RNA synthesis catalyzed by EV-71 RdRp using a template with a single GS-443902 or GS-646939 residue in the template at position 11.** ***A***, RNA primer/template with a template-embedded GS-443902 (*Template "GS-443902"*, *middle*) or GS-646939 (*Template "GS-646939"*, *right*) at position 11; the corresponding primer/template with adenosine (*Template "A"*) at this position is shown on the left. G5 indicates incorporation of [α-$^{32}$P]-GTP at position 5. ***B***, Migration pattern of the products of RNA synthesis catalyzed by EV-71 RdRp. $MgCl_2$, [α-$^{32}$P]-GTP, and ATP were provided to the reaction to support RNA synthesis up to position 10. Increasing concentrations of UTP were supplemented to the reactions to monitor incorporation opposite adenosine, GS-443902, or GS646939 at position 11, and templated adenosines from position 12 to 20. For Template "GS-443902", intermediate products are observed at position 10. For Template "GS-646939", additional sites of inhibition are indicated by the formation of intermediate products at positions 9 and 14. Product formation at and beyond the asterisk indicates RNA products that are likely a result of sequence-dependent slippage events. A 5′-$^{32}$P-labeled 4-nt primer serves as a size marker ***C***, Quantification of ***B*** (*top*) where the sum of RNA products generated beyond position 10 was divided by the total signal in the lane and normalized as a percentage, fold-inhibition resulting from a templated GS-443902 and GS-646939 (*bottom*). To account for template-dependent differences in activity, product fraction was normalized as a percentage to the product fraction observed at 1000 μM UTP for that template.

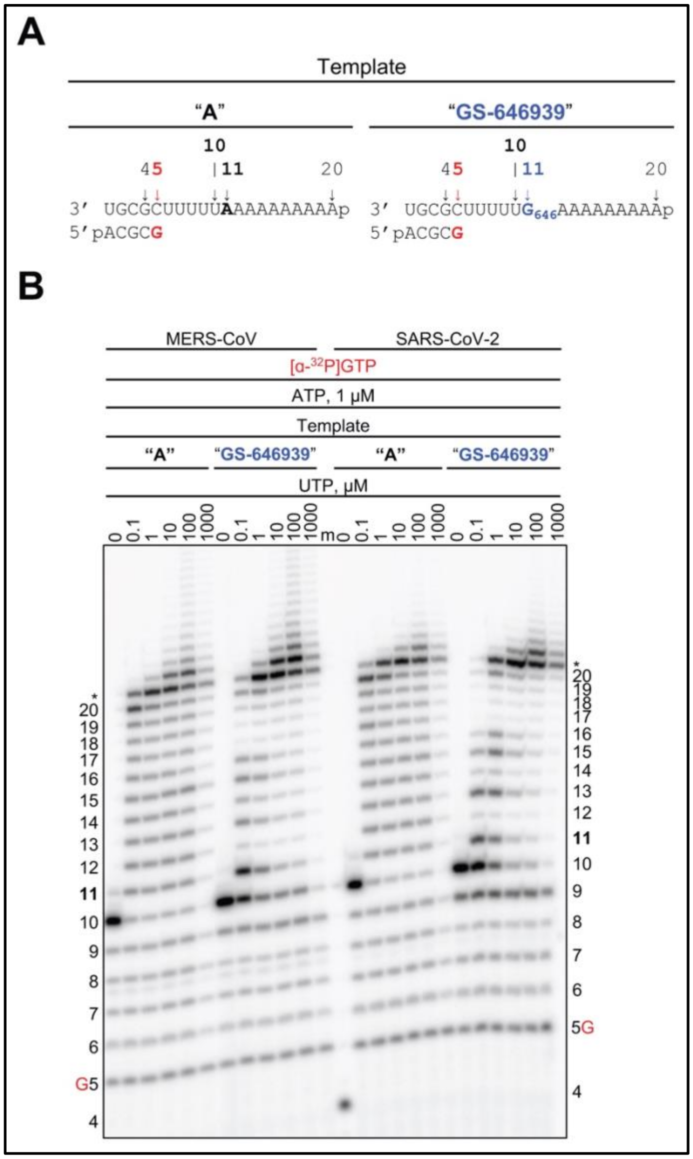


**Figure S9: RNA synthesis catalyzed by MERS-CoV and SARS-CoV-2 RdRp using a template with a single GS-646939 residue embedded at position 11.** ***A***, RNA primer/template with template-embedded GS-646939 (*Template "GS-646939", right*); the corresponding primer/template with adenosine (*Template "A"*) at this position is shown on the left. G5 indicates incorporation of [α-$^{32}$P]-GTP at position 5. ***B***, Migration pattern of the products of RNA synthesis catalyzed by MERS-CoV and SARS-CoV-2 RdRp. $MgCl_2$, [α-$^{32}$P]-GTP, and ATP were provided to the reaction to support RNA synthesis up to position 10. Increasing concentrations of UTP were supplemented to the reactions to monitor incorporation opposite an embedded AMP, or GS-646939, at position 11, and templated adenosines from position 12 to 20. A 5′-$^{32}$P-labeled 4-nt primer serves as a size marker.
